# GABA-Glutamate supramammillary neurons control theta and gamma oscillations in the dentate gyrus during paradoxical (REM) sleep

**DOI:** 10.1101/584862

**Authors:** Francesca Billwiller, Laura Castillo, Heba Elseedy, Anton Ivanovich Ivanov, Jennyfer Scapula, Antoine Ghestem, Julien Carponcy, Paul Antoine Libourel, Hélène Bras, Nabila ElSayed Abdelmeguid, Esther Krook-Magnuson, Ivan Soltesz, Christophe Bernard, Pierre-Hervé Luppi, Monique Esclapez

**Affiliations:** UMR 5292 CNRS/U1028 INSERM, Centre de Recherche en Neurosciences de Lyon (CRNL), Université Claude Bernard Lyon I, LYON Cedex 08, France; Aix Marseille Univ, INSERM, INS, Institut de Neurosciences de Systèmes, Marseille, France; Aix Marseille Univ, CNRS, INT, Institut de Neurosciences Timone, Marseille, France; Zoology Department, Faculty of Science, Alexandria University, Alexandria, Egypt; Department of Neuroscience, University of Minnesota, Minneapolis, MN 55455, USA; Department of Neurosurgery, Stanford University, USA

## Abstract

Several studies suggest that neurons from the lateral region of the SuM (SuML) innervating the dorsal dentate gyrus (DG) display a dual GABAergic and glutamatergic transmission and are specifically activated during paradoxical (REM) sleep (PS). The objective of the present study is to fully characterize the anatomical, neurochemical and electrophysiological properties of the SuML-DG projection neurons and to determine how they control DG oscillations and neuronal activation during PS and other vigilance states. For this purpose, we combine structural connectivity techniques using neurotropic viral vectors (rabies virus, AAV), neurochemical anatomy (immunohistochemistry, in situ hybridization) and imaging (light, electron and confocal microscopy) with *in vitro* (patch clamp) and *in vivo* (LFP, EEG) optogenetic and electrophysiological recordings performed in transgenic VGLUT2-cre male mice. At the cellular level, we show that the SuML-DG neurons co-release GABA and glutamate on dentate granule cells and increase the activity of a subset of DG granule cells. At the network level, we show that activation of the SuML-DG pathway increases theta power and frequency during PS as well as gamma power during PS and waking in the DG. At the behavioral level, we show that the activation of this pathway does not change animal behavior during PS, induces awakening during slow wave sleep and increases motor activity during waking. These results suggest that the SuML-DG pathway is capable of supporting the increase of theta and gamma power in the DG observed during PS and plays an important modulatory role of DG network activity during this state.

**Significant statement:** An increase of theta and gamma power in the dentate gyrus (DG) is an hallmark of paradoxical (REM) sleep (PS) and is suggested to promote learning and memory consolidation by synchronizing hippocampal networks and increasing its outputs to cortical targets. However the neuronal networks involved in such control of DG activity during PS are poorly understood. The present study identifies a population of GABA/Glutamate neurons in the lateral supramammllary nucleus (SuML) innervating the DG that could support such control during PS. Indeed, we show that activation of these SuML-DG projections increase theta power and frequency as well as gamma power in the DG specifically during PS and modulate activity of a subset of DG granule cells.

## Introduction

The supramammillary nucleus (SuM) is a thin structure overlying the mammillary bodies in the hypothalamus that provides substantial projections to many regions of the limbic system including the hippocampal formation (for review see Pan and McNaughton 2004, Vertes 2015). The SuM is composed of several populations of neurons that differ in their neurochemical content and the specificity of their connections (Swanson et al., 1982; Gonzalo-Ruiz et al., 1992; Leranth and Kiss, 1996; Kocsis et al., 2003; Borhegyi and Leranth, 1997; Haglund et al., 1984). In rats, many studies have described the projections from SuM neurons to the hippocampus (Segal & Landis, 1974; Pasquier & Reinoso-Suarez, 1976, 1978; Wyss et al., 1979; Haglund et al., 1984; Saper et al., 1985; Vertes, 1992; Magloczky et al., 1994; Nitsch and Leranth 1996; Vertes & McKenna, 2000). We (Soussi et al., 2010) and others (Boulland et al., 2009) demonstrated in rats that SuM neurons from the most medial part of the SuM (SuMM; Paxinos and Watson, 1998), referred as SuMm by Swanson (1998), or SuMp by Pan and McNaughton (2004) innervating the inner molecular layer of the ventral dentate gyrus (DG) and CA2 pyramidal cells are glutamatergic (Soussi et al., 2010). In contrast, SuM neurons located in the lateral two-third region of the SuM (SuML), referred as SuMg by Pan and McNaughton (2004), innervate the supragranular layer of the dorsal DG (dDG) and to a lesser extend the ventral DG (vDG) and display a unique dual glutamatergic and GABAergic neurotransmitter phenotype (Boulland et al., 2009; Soussi et al., 2010). Indeed, these SuML neurons and their projections to the dDG co-express markers for both glutamatergic (vesicular glutamate transporter 2; VGLUT2) and GABAergic (glutamate decarboxylase 65, GAD65 and vesicular GABA transporter, VGAT) neurotransmission and establish asymmetric and symmetric synapses on DG cells (Soussi et al., 2010). The connectivity and neurochemical properties of these different SuM-hippocampal pathways suggest that they may contribute differently to hippocampal dependent functions. Indeed, it has been shown that the SuM controls hippocampal theta rhythm (Kocsis and Vertes 1994, Vertes and Kocsis 1997, Kocsis and Kaminski 2006) and is involved in emotional learning and memory (Pasquier and Reinoso-Suarez 1976, Richmond et al 1999, Pan and McNaughton 2002, Santin et al 2003, Pan and McNaughton 2004, Shahidi et al. al., 2004). By combining retrograde tracing, neurotoxic lesion and FOS immunostaining, it was recently shown that SuML GABA/Glutamate neurons are responsible for the activation of dDG granule cells during paradoxical sleep (PS) (Renouard et al., 2015; Billwiller et al., 2016). Such activation may play a key role in the previously reported beneficial effect of PS on learning and memory (Maquet et al., 2000). In addition, Pedersen and colleagues (2017), by using a chemogenetic approach in transgenic mice, showed that SuM neurons containing only VGLUT2 but not those containing VGAT and VGLUT2 play a crucial role in waking. Altogether these results suggest that glutamatergic neurons from the SuM known to innervate several limbic regions including the CA2/CA3a hippocampal region and the vDG are involved in normal wakefulness whereas the GABA/Glutamate SuML neurons projecting to the dDG could be instrumental for PS function.

In this study we investigate how the SuML-dDG pathway control dDG neurons activity during PS and other vigilance states. For this purpose we combine innovative structural connectivity techniques using neurotropic viral vectors (rabies virus, AAV), neurochemical anatomy (immunohistochemistry, *in situ* hybridization) and imaging (light, electron and confocal microscopy) with *in vitro* (patch clamp) and *in vivo* (LFP, EEG) optogenetic and electrophysiological recordings performed in transgenic VGLUT2-cre mice in order: 1) to fully characterize the anatomical, neurochemical and electrophysiological properties of the SuML-dDG projection neurons in our mice trangenic model and 2) to determine the influences of this specific pathway on behaviors and associated oscillatory activities characterizing the different vigilance states as well as DG neuron activity.

## Materials and methods

### Animals

For this study, we used 26 transgenic “VGLUT2-cre” adult male mice (25-30 g; aged 10-11 weeks) obtained by mating male and female homozygote Vglut2-ires-Cre mice from Jackson Laboratories (Strain name Slc17a6^tm2(cre)Lowl^/J, stock #016963). All mice were bred in-house and maintained in standard cages, with food and water *ad libitum*, in a temperature- and humidity-controlled room under a 12 hr light/12 hr dark cycle. All the surgical and experimental procedures were performed according to the National Institutes of Health guidelines and the European communities Council Directive of 86/609/EEC and were approved by the University of Aix-Marseille and Lyon University Chancellor’s Animal Research Committees.

### Vectors

#### Rabies virus

Four VGLUT2-cre mice underwent stereotaxic injection of rabies virus (RV) within the inner molecular layer of the dDG in order to obtain golgi-like retrograde-labeling of SuM neurons projecting to this region. The strain of RV used was the Challenge Virus Standard (CVS, 4.10^7^ plaque-forming units/mL) (Bras et al. 2008; Ugolini 2010; Coulon et al. 2011). Only vaccinated personnels conducted these experiments at the appropriate biosafety containment level until the sacrifice of the animals as described below.

These animals were used to determine the neurotransmiter phenotype of SuM neurons projecting to dDG at the mRNA level, combining immunohistofluorescent detection of RV with fluorescent *in situ* hybridization detection of VGAT and VGLUT2 mRNAs.

#### Adeno-Associated Virus (AAV) Double floxed Inverted ORF (DIO) vectors

Thirteen VGLUT2-cre mice underwent stereotaxic injection of the cre-dependent viral vectors: AAV5-EF1a-DIO-EYFP (3.5 10^12^ virus molecules / ml; UNC Gene Therapy Center Vector Core; Dr Deisseroth) into the SuML. These VGLUT2-EYFP mice expressing the Yellow Fluorescent Protein (YFP) in VGLUT2 SuM neurons and their axon terminals were used 1) to determine the neurotransmitter phenotype of SuM neurons innervating the dDG at the protein level, by simultaneous immunohistofluorescent detection of EYFP labeled axon fibers and terminals, VGAT and VGLUT2 in dDG (n=3); 2) to determine the synaptic profile of these EYFP labeled axon terminals at the electron microscopy level (n=3); 3) as control animals (n=3) for *in vitro* optogenetic stimulation and patch clamp electrophysiological recordings; 4) as control animals (n=4) for *in vivo* optogenetic stimulation and electrophysiological recordings followed by cFos immunolabeling.

Nine VGLUT2-cre mice underwent stereotaxic injection of the cre-dependent viral vectors: AAV5-EF1a-DIO-hChR2(H134R)-EYFP (3.2 10^12^ virus molecules / ml; UNC Gene Therapy Center Vector Core; Dr. Deisseroth) into the SuML. These VGLUT2- ChR2 mice expressing the excitatory opsin, channelrhodopsine 2 (ChR2) and the reporter protein EYFP in VGLUT2 SuM neurons and their axon terminals were used for 1) *in vitro* optogenetic stimulation and patch clamp electrophysiological recordings experiments (n=5); 2) *in vivo* optogenetic stimulation and electrophysiological recordings followed by cFos immunolabeling (n=4).

### Surgery

Mice were anesthetized by an intraperitoneal injection (i.p.) of Ketamin (50mg/kg) / Xylazine (5mg/kg) solution. If necessary, this anesthesia was repeated during the surgery. Animals were then secured in a stereotaxic frame (David Kopf instruments). The body temperature of mice was monitored and maintained at about 37°C during the entire procedure by means of an anal probe and heating blanket respectively. The head was shaved and sanitized with Betadine and 0.9% NaCl. Local anesthesia was performed by infiltration of the scalp with xylocaine (lidocaine hydrochloride 0,5%), and an ophthalmic gel was placed on the eyes to avoid drying. After scalp incision, holes were drilled in the skull, with antero-posterior (AP), medio-lateral (ML) and dorso-ventral (DV) coordinates based on Paxinos & Franklin’s atlas (2005).

#### RV and AAV viral vector injections

The RV (200 nl) was pressure-injected unilaterally (n=2) or bilaterally (n=2) within the dDG inner molecular layer of VGLUT2-cre mice according to the following coordinates: AP = −2; ML = +/-1.5; DV = −1.7. Injections were performed by using a 33-gauge Hamilton syringe connected to a Micro4 injection pump system (World Precision Instruments). After completion of the injection procedures, the syringe was removed and the skin was sutured. Animals were treated with local anesthetic, returned to their cages kept at the appropriate biosafety containment level for a survival period of 38h to observe an optimal RV retrograde labeling for the dendritic arbor of SuM neurons. AAV5-EF1a-DIO viral vectors (500nl) were injected bilaterally within the SuML of VGLUT2-cre mice at the following coordinates: AP = −2.7; LM = +/-1.25; DV = −4.8 using an 11°angle to avoid the high vascularization at the midline.

#### Optrode and electrode implantation

After AAV injections, VGLUT2-mice were implanted with an optrode in the left dDG (AP = −2; ML = −1.1; DV = −1.7) for optic stimulation of axon terminals originating from transfected SuML neurons and local field potential (LFP) recordings from the DG. The optrode consisted of an optic fiber (250 μm ∅, 0.39 NA; Thorlabs SAS) and the LFP electrode. The LFP electrodes consisted of two 45 µm diameter tungsten wires (California Fire Wire Company), twisted and glued together to form a rigid and solid structure, with the 2 ends of the wires separated by 100 µm from each other. Two small screws (1 mm in diameter, Plastics One) each soldered to a wire were fixed on the skull. The first screw was fixed in the parietal part of the skull for EEG recording; the second screw, at the level of cerebellum as reference electrode. Two wire electrodes were inserted into the neck muscles for bipolar EMG recordings. All leads were connected to a miniature plug (Plastics One) that was cemented on the skull.

After completion of the surgery the skin was sutured. Mice were treated with local anesthetic, and an intramuscular injection of antibiotic (Baytril, 5mg/kg) to prevent any risk of infection. Mice were monitored until waking and replaced in their home cages for a survival period of 3 weeks.

### Histology

#### Tissue preparation for light microscopy

Animals were deeply anesthetized with ketamine and xylazine and transcardially perfused with 4% paraformaldehyde (PFA) in 0.12 M sodium phosphate buffer, pH 7.4 (PB). After perfusion, the brains were removed from the skull, post-fixed in the same fixative for 1 h at room temperature (RT), rinsed in PB, cryoprotected in 20% sucrose overnight, frozen on dry ice and sectioned coronally at 40 µm with a cryostat (Microm). The sections were rinsed in PB, collected sequentially in tubes containing an ethylene glycol-based cryoprotective solution and stored at −20°C until histological processing. One of every ten sections was stained with cresyl violet to determine the general histological characteristics of the tissue throughout the rostro-caudal extent of the brain. Selected sections were processed for 1) simultaneous detection of VGLUT2 mRNA, VGAT mRNA and RV; 2) simultaneous immunohistofluorescent detection of EYFP, VGLUT2 and GAD65 or VGAT; 3) immunohistochemical detection of cFos.

#### Simultaneous detection of VGLUT2 mRNA, VGAT mRNA and RV combining fluorescent in situ hybridization (RNAscope technology) and immonhistofluorescent methods performed in VGLUT2-cre mice injected with RV

Selected sections at the level of the SuM were first treated with 1% H2O2 rinsed in PB mounted on SuperFrost Plus slides (Fisher Scientific) and air dried at RT. They were then processed for fluorescent RNAscope *in situ* hybridization according to the manufacter’s protocol (Advanced Cell Diagnostics). Briefly sections were treated with 100% ethanol and protease III for 30 min at 40°C. They were incubated in a solution containing both RNAscope® Probe – Mm-S1c17a6 for detection of VGLUT2 mRNA and Mm-S1c32a1-C3 for detection of VGAT mRNA. After hybridization, sections were then processed for visualization using the RNA-scope Multiplex Fluorescent reagent Kit v2 (Advanced Cell Diagnostics) and the Tyramide Signal Amplification (TSA™) Plus Cyanine 3 and TSA Plus Cyanine 5 systems (Perkin Elmer).

After the RNAscope assay, sections were rinsed in 0.02 M potassium phosphate-buffered saline (KPBS, pH 7.2-7.4) and processed for immnohistofluorescent detection of the RV using a Mouse On Mouse kit (MOM, Vector Laboratory) to eliminate non specific labeling due to use of mouse monoclonal antibody on mouse tissue. Sections were incubated for 1 h at RT in MOM mouse IgG blocking reagent diluted in KPBS containing 0.3% Triton X-100. Sections were rinsed twice with KPBS for 5 min and pre-incubated in MOM diluent for 15 min. Sections were then incubated overnight at RT in a solution containing the mouse monoclonal antibody directed against RV (1:3000; Raux et al., 1997 Kindling provided By Dr. Patrice Coulon), diluted in MOM diluent. After several rinses in KPBS, they were incubated for 2 h in Alexa488-conjugated donkey anti-mouse (1:200; Invitrogen) diluted in MOM diluent. After several rinses in KPBS, all sections were coverslipped with Fluoromount (Electron Microscopy Sciences). The specimens were analyzed with a Zeiss laser-scanning confocal microscope.

#### Quantification of co-localizing RV, VGLUT2 and VGAT mRNAs

Quantitative analysis was conducted to evaluate the extent of SuM neurons with direct projections to the dDG that co-express VGLUT2 and VGAT mRNAs. For this purpose, the number of triple neurons was determined for each animal (n=4), from 3 sections (120 µm apart from each other) across the antero-posterior extent of the SuM. For each section, an image of the entire SuM region was obtained from a single confocal slice using the Tile Scan function with a 20x objective and sequential acquisition of the different wavelength channels to avoid fluorescent talk with ZEN software (Zeiss). The analysis was then performed with Neurolucida software (version 7, mbfBioscience) as follows: for each confocal image, all RV-labeled neurons were identified on the green channels and examined for colocalization of VGLUT2 mRNA in the blue channel and/or VGAT mRNA in the red one. Triple- double- and single-labeled neurons were tagged differently and counted by the software. A total of 380 RV-labeled neurons were analyzed.

#### Simultaneous immunohistofluorescent detection of EYFP, VGLUT2 and GAD65 or VGAT performed in VGLUT2-EYFP mice

Selected sections at the level of the dDG were processed using the MOM kit as described above. Sections were incubated overnight at RT in a solution containing rabbit anti-GFP (1:2000, Invitrogen), guinea pig anti-VGLUT2 (1:5000, Millipore) and mouse anti-GAD65 (1:100, Millipore) or mouse anti-GFP (1:100, Invitrogen), guinea pig anti-VGLUT2 (1:5000, Millipore) and rabbit anti-VGAT (1:1000, Synaptic System) diluted in MOM diluent. After several rinses in KPBS, they were incubated for 2 h in Alexa488-conjugated donkey anti-rabbit IgG (1:200; Invitrogen), Cy5-conjugated donkey anti-guinea pig (1:100; Jackson ImmunoResearch Laboratories, Inc.), and Cy3-conjugated donkey anti-mouse (1:100; Jackson ImmunoResearch Laboratories, Inc.) or Alexa488-conjugated donkey anti-mouse IgG (1:200; Invitrogen), Cy5-conjugated donkey anti-guinea pig (1:100; Jackson ImmunoResearch Laboratories, Inc.), and Cy3-conjugated donkey anti-rabbit (1:100; Jackson ImmunoResearch Laboratories, Inc.) diluted in MOM diluent. After several rinses in KPBS, all sections were then mounted on superfrost-coated slides, dried overnight at RT and coverslipped with Fluoromount. The specimens were analyzed with confocal microscope (Zeiss).

#### Quantification of co-localizing GFP, VGLUT2 and VGAT

Two quantification protocols were used in order to evaluate the extent of the different neurochemical phenotypes of axon terminals from the SuM innervating the dDG. In the first protocol previously described (Soussi et al., 2015), the densities of VGLUT2/VGAT and VGLUT2 only labeled terminals were assessed by the quantification of immunolabeling for VGLUT2/VGAT and VGLUT2 only, respectively. These analyses were performed on 4 sections for each mouse. Single optical confocal images were acquired with Zeiss LSM 510 laser-scanning microscope and analyzed with the software provided by the microscope manufacturer (LSM 510 Zen, Zeiss). All images were acquired from the suprapyramidal and infrapyramidal blades of the dDG, using identical parameters. The percentages of VGLUT2 labeled terminals containing VGAT were estimated by the the Manders’ coefficient (proportion of pixels for VGLUT2 also positive for VGAT) obtained with the JACoP co-localization Plugin for Image J, in the region of interest (ROI) which included granule cell layer (GCL) and the narrow zone superficial to the granule cells defined as the supragranular layer (SGL) following recommendations from Bolte & Cordelières (2006). For each channel, an identical bottom threshold was applied throughout the analyses, and only the pixels with a value above this threshold were counted. When a pixel had a value above the threshold in both channels, it was counted as double positive. The size and the shape of the ROI was the same for each confocal image. The average % of co-localization was calculated for each blade of the DG for each mouse.

In the second quantification protocol, we determined for each mouse (n=4), the relative percentages of triple- and double-labeled boutons for the GFP anterograde tracer and VGAT and ⁄ or VGLUT2 in the SGL and infragranular blade of the dDG from several z-stacks of 10 confocal slices, acquired with the 100X objective and a numerical zoom 8, in three different sections. The analysis was performed as previously described (Persson et al., 2006; Soussi et al., 2010) and following recommendations from Bolte & Cordelières (2006). For each z-stack, the confocal images obtained from separate wavelength channels (green, red and blue) were displayed side by side on the computer screen together with the images corresponding to colocalized pixels within each optical slice of the z-stack obtained with the colocalization highlighter plugin in ImageJ. The GFP-labeled boutons were identified in the green channel within a probe volume defined by the size of the confocal slice (19.38 μm · 19.38 μm) and the height of the z-stack (2 μm). Each bouton was examined for colocalization through the individual optical slices of the z-stack. Single-, double- and triple-labeled boutons were counted using the Cell Counter plugin in Image J. The total number of GFP-labeled terminals analyzed in the two regions of interest was 400.

#### Immunohistochemical detection of cFos performed in VGLUT2-EYFP (n=4) and VGLUT2-ChR2 (n=4) mice

Selected sections at the level of the DG were processed for immunohistochemistry according to previously described protocol (Esclapez et al. 1994). Sections were pre-treated for 30 min in 1 % H2O2, rinsed in PB and KPBS, preincubated for 1 h in 3 % normal goat serum (NGS, Vector Laboratories) diluted in KPBS containing 0.3 % Triton X-100 and incubated overnight at RT in cFos rabbit polyclonal antiserum (1:20,000; Calbiochem) diluted in KPBS containing 1 % NGS and 0.3 % Triton X-100. After several rinses in KPBS, sections were incubated for 1 h at RT in biotinylated goat anti-rabbit immunoglobulin G (IgG; Vector Laboratories) diluted 1:200 in KPBS containing 3 % NGS and then for 1 h at RT in an avidin-biotin-peroxidase complex solution prepared in KPBS according to the manufacturer’s recommendations (Vectastain ABC kit, Vector Laboratories). Sections from VGLUT2-EYFP and VGLUT2-ChR2 mice were processed In parallel and for the same period of time (15 min) in 3.3’-diaminobenzidine tetrahydrochloride (DAB, Sigma fast tablets; Sigma), rinsed in KPBS, mounted onto Superfrost Plus slides, dehydrated and coverslipped with Permount.

#### Quantification of cFos immunolabeled neurons

The number of cFos labeled neurons was calculated in the DG GCL of the right and left (ipsilateral to the optic stimulation) hemispheres in VGLUT2-EYFP control (n=4), and VGLUT2-ChR2 (n=4) mice. These analyses were performed using a computer-assisted system connected to a Nikon 90i microscope and the Neurolucida software (MicroBrightField). A total of 4 sections (400 µm apart from each other) surrounding the optrode site were analyzed for each animal. In each section the GCL was delineated and all neurons labeled for cFos were plotted. The software calculated the total number of labeled neurons in each hemisphere for each animal. The average total number of labeled neurons / hemisphere ± SEM was calculated for each group of control VGLUT2-EYFP and VGLUT2-ChR2 mice. Statistical analysis was performed by Statview software using Wilcoxon Rank Sum Test.

#### Tissue preparation for electron microscopy

Three VGLUT2-EYFP mice were perfused intracardially with a fixative solution containing 4% PFA and 0.1% glutaraldehyde in 0.12 m PB. After perfusion, the brain was removed from the skull, post-fixed in the same fixative overnight at 4°C and rinsed in PB for 1.5 h. Blocks of the forebrain were sectioned coronally at 60 µm with a vibratome. Pre-embedding immunolabeling for GFP sections at the level of dDG were pre-treated for 15 min in 1% sodium borohydride prepared in PB and rinsed for 30 min in PB and 3 × 30 min in KPBS. Sections were incubated for 1h in normal goat serum diluted in 0.02M KPBS, then incubated overnight in primary antibody rabbit anti-GFP (1:2000) diluted in KPBS containing normal goat serum at RT. On the following day sections were rinsed for 1.5 h in 0.02M KPBS then incubated for 1h in the secondary antibody goat anti-rabbit (1:200) diluted in KPBS containing normal goat serum. After rinsing in KPBS for 1.5 h, sections were incubated for 1h in an avidin-biotinylated-peroxidase complex (ABC Elite; Vector Laboratories) prepared in KPBS. After 3 × 30 min rinses in KPBS, sections were incubated for 12 min in 3.3’-diaminobenzidine tetrahydrochloride and 0.01% H2O2, rinses in KPBS, post-fixed in 2% PFA and 2.5% Glutaraldahyde diluted in PB for 3 h, then washed for 1.5 h in PB. After all these steps, sections were treated with 1% osmium tetroxide in PB for 45 min, dehydrated in ethanol, flat embedded in Durcupan resin and polymerized at 56°C for 24 h (Zhang & Houser, 1999). Labeled regions of the DG that contained the molecular and granule cell layers were trimmed from the sections, re-embedded on capsules filled with polymerized Durcupan and further polymerized at 56°C for an additional 24 h. Ultrathin sections from the most superficial face of the blocks were cut on an ultramicrotome. Serial sections were picked up on nickel mesh grids and stained with uranyl acetate and lead citrate. Sections were examined and photographed with a JEOL electron microscope.

### *In vitro* electrophysiology: optic stimulation and patch clamp recordings

#### Hippocampal slice preparation

VGLUT2-ChR2-EYFP mice (n=5) and VGLUT2-EYFP mice (n=3) were decapitated under isofluorane anesthesia. Brains were quickly removed and placed into an ice-cold (4°C) cutting solution containing (in mM): 140 potassium gluconate, 10 HEPES, 15 sodium gluconate, 0.2 EGTA, 4 NaCl (pH 7.2). Coronal slices were cut (350 µm) using a vibratome (Leica Microsystem). In order to increase cell survival over time in slices from 10 to 11 week-old adult, slices were incubated first in a solution containing (in mM): 110 choline chloride; 2.5 KCl; 1.25 NaH_2_PO_4_; 10 MgCl_2_, 0.5 CaCl_2_; 25 NaHCO_3_; 10 glucose, 5 sodium pyruvate, for 15 min at 20-23°C. Then, they were transferred to a holding chamber containing an artificial cerebrospinal fluid (ACSF) composed of (in mM) 126 NaCl, 3.5 KCl, 1.2 NaH_2_PO_4_, 1.3 MgCl_2_, 2 CaCl_2_, 25 NaHCO_3_, 10 D-glucose (pH = 7.3-7.4) at RT for at least 1 h before recording. The two last solutions were saturated with 95% O_2_ and 5% CO_2_.

#### Whole-cell voltage-clamp recordings

Slices were submerged in a low-volume recording chamber and continuously superfused with 32-34°C ACSF at 5 ml/min perfusion rate. For each mouse, four slices containing the dDG were selected for patch clamp recordings. DG neurons were visualized by infrared video microscopy using an upright microscope (SliceScope, Scentifica Ltd). Patch pipettes were pulled from borosilicate glass tubing (1.5 mm outer diameter, 0.5 mm wall thickness) and filled with an intracellular solution containing (in mM) 20 CsCl, 115 CsGlu, 10 HEPES, 1.1 EGTA, 4 MgATP, 10 Na phosphocreatine and 0.4 Na_2_GTP as well as 0.2% biocytin for *post-hoc* morphological identification of the recorded neuron (see below). The pipette resistance was 4–6 MΩ. Recordings were performed in the Apex, upper and lower blades of the dDG. Signals were fed to a Multiclamp 700A (Molecular Devices), digitized (10 kHz) with a DigiData 1550 (Molecular Devices) interface to a personal computer and analysed with ClampFit software (Molecular Devices). Optical stimulation of ChR2-expressing axon terminals was performed by pulses of 470 nm blue light delivered by a LED (pE-2, CoolLED) through a 40X objective attached to microscope (SliceScope, Scientifika Ltd). Stimulations consisted of paired 5 ms pulses (500 ms between pulses, every 30 s). For VGLUT2-ChR2 mice, the light intensity corresponded to 20-30% (1.6-2.5 mW) of the LED maximum power (7.5 mW) and for the VGLUT2-EYFP control mice, it varied between 20 to 90% (6.9 mW) of LED maximum power. Postsynaptic current (PSC) responses to optic stimulations were recorded at different holding potentials ranging from −70 mV (close to reversal potential of GABA-A currents but far below the reversal potential of glutamate receptor-mediated currents) to +10mV (close to reversal potential of glutamate receptor-mediated currents but far above the reversal potential of GABA-A-mediated currents). Pharmacological characterization of inhibitory PSCs (IPSCs) and excitatory PSCs (EPSCs) was achieved using antagonists of GABA-A, AMPA and NMDA receptors. We used Gabazine or bicuculline (antagonist of GABA-A receptors, 10 µM), D-AP5 (antagonist of NMDA receptors, 40 µM) and NBQX (antagonist of AMPA and Kaïnate receptors, 10 µM). The co-release of glutamate and GABA was further demonstrated in several DG granule cells (n=5), by recording first light stimulated PSC at −70 mV (Figure 4K). At this holding potential the recorded currents were essentially generated by glutamate receptors. Then the glutamate component was abolished by NBQX and D-AP5, and the GABA component was revealed at +10 mV holding potential. This PSC was completely inhibited by bicuculline application that confirmed the GABA receptor origin of these remaining currents (see also Figure 4J).

#### Double immunohistofluorescent labeling for Biocytin and GFP

After recordings slices were processed for simultaneous detection of the biocytin-filled neurons and GFP-labeled axon fibers and terminals in order to identify the recorded cells and evaluate the efficiency of the transfection, respectively. Slices were fixed overnight at 4°C in a solution containing 4% PFA in PB. Then they were rinsed in PB, cryoprotected in 20% sucrose and quickly frozen on dry ice. After several rinses in KPBS, slices were incubated in a solution containing normal donkey serum (NDS, 1:30; Vector Laboratory) diluted in KPBS with 0.3% Triton-X100, for 2 h at RT. They were incubated in a solution containing goat anti-biotin (1:200) and rabbit anti-GFP (1:2000), diluted in KPBS containing 0.3% Triton-X100 and NDS (1:100), overnight at RT. After several rinses in KPBS, slices were incubated for 2 h in Alexa488-conjugated donkey anti-goat IgG (1:200); Invitrogen), and Cy3-donkey anti-rabbit IgG (1:100) diluted in KPBS with 0.3% Triton-X100. After rinses in KPBS, slices were mounted on slides and coverslipped with Fluoromount. The specimens were analyzed with a fluorescence microscope (Nikon 50i) or confocal microscope (Zeiss).

### *In vivo* electrophysiology: optic stimulation, LFP and EEG recordings

All mice were placed for 7 days in a recording box in order to get them used to the recording conditions. The recording box was ventilated, as well as electrically and sound isolated. The temperature was regulated at 21°C, and a 12 h light/12 h dark cycle imposed. Mice were accustomed to the cable connecting them to the recording device. The recording cable connected the micro-connector implanted on the head of the animal to a collector, which ensured the continuity of the recorded signals without hindering the movements of the mouse. At the end of this habituation, the control recordings begun. EEG and EMG recordings were digitized at 1 kHz, amplified 5000 times with a 16 channels amplifier (A-M System) and collected on a computer via a CED interface using Spike 2 software (Cambridge Electronic Design). The signal was band-pass filtered online between 1 and 300 Hz for EEG, and between 10 and 100 Hz for EMG. The 50 Hz signal was removed with a notch filter. The EEG and LFP signals were acquired by monopolar derivation (differential between the recording electrode and the reference electrode located above the cerebellum). The EMG bipolar signals were calculated by measuring the differential between the two EMG electrodes. Mice were recorded for 24 h of baseline followed by optogenetic manipulation.

#### In vivo optogenetic stimulation

Optical stimulations were delivered via a patch cable connected to a 100 mW 473-diode (Laserglow). Stimulations were performed during 4 days: in the first 3 days, mice were stimulated during one specific vigilance state par day: waking (WK), slow wave sleep (SWS) or PS. Each day, stimulations were delivered during the same circadian period (10 AM–2 PM). Stimulations were applied 10 s after the occurrence of a stable WK, SWS or PS event as detected by real time observation by the experimenter. For WK and SWS, stimulations were spaced apart by at least 1 min and for PS, by at least 15 s.

Blue exciting stimulations consisted of 10-s trains of 10-ms pulses at 20 Hz. Light power at the fiber tip was 10 mW.

The 4^th^ day of experiments, 4 VGLUT2-EYFP and 4 VGLUT2-ChR2 animals were stimulated for 15 min at 20 Hz (10-ms pulses). All mice were euthanized 90 min after the beginning of the stimulation by transcardiac perfusion of 4% PFA. Brain tissues were processed for immunohistochemical detection of cFos expression (see above).

#### Analysis of the sleep wake states

Polysomnographic recordings were visually scored by 5 s epochs for WK, SWS and PS as previously described (Sapin et al. 2009). Hypnograms were obtained by using a custom Matlab script. For each animal, the number of awakenings during SWS and PS optogenetic stimulations was counted and expressed as percentage of the total number of stimulations.

#### LFP and EEG analysis

LFP and EEG signals were analyzed using a custom Matlab script using the Chronux toolbox. The time-frequency spectrograms were computed with the same toolbox and expressed in arbitrary units. The mean power spectral density in the 10 s before the stimulation was compared to that in the 10 s during the stimulation (sliding window: 1 s), in order to obtain a mean spectral power ratio (PR) ± SEM. The frequency spectra were grouped into frequency bands commonly used Delta: 1-4 Hz, Theta: 6-12 Hz, Sigma: 12-14 Hz, Beta: 15-30 Hz, Gamma: 30-100 Hz. Power spectral values at 20 Hz and its harmonics were excluded from the analysis.

To analyze the evolution of LFP and EEG theta and gamma bands during optogenetic stimulation, the mean PR of these spectral bands and the respective 95% confidence intervals were calculated from 10 s before to 10 s after the photostimulation. In order to compute these intervals, we used a bootstrap procedure, which allows creating artificial groups from the original data, with replacement. The mean of each artificial group derived from the original data was then computed. This operation was repeated 10000 times and the 95% confidence interval was the 5^th^ and the 95^th^ percentile of the means of the randomly constructed samples.

Finally, during WK and PS the peak of theta frequency (6-12 Hz) in the 10 s before and during the optogenetic stimulation was identified in all animals.

Analysis of variance (Mann-Whitney test) was performed on the percentage of awakenings, the mean spectral power ratios and the theta frequency peaks. These statistics were performed using Statview software (StatView Inc, Nestbit, NS).

#### EMG analysis

EMG signals during WK were analyzed using a custom Matlab script. The mean EMG value in the 10 s before the stimulation was compared to that in the 10 s during the stimulation, in order to obtain a mean EMG ratio ± SEM. In the 2 groups of animals we performed a sequential analysis per 0.5 s on the mean of the absolute EMG values from 20 seconds before to 10 seconds after the photostimulation. The respective bootstrap 95% interval (computed as described above) was calculated for each sequential value. Further, analysis of variance (Mann-Whitney test) was performed on the EMG mean by using Statview software.

## Results

### Distribution of VGAT and/or VGLUT2 mRNA-containing neurons in the SuM of VGLUT2-Cre transgenic mice

Three populations of VGAT and/or VGLUT2 mRNA-containing neurons were observed at all antero-posterior levels of the SuM. These included a population of large neurons co-expressing VGAT and VGLUT2 mRNA (Figure 1 A, D, arrows); and two populations of smaller neurons containing either VGAT mRNA (Figure 1 A, D, arrowheads) or VGLUT2 mRNA (Figure 1 A, D, blue). The distribution as well as the density of these populations differed significantly within the SuM. Almost all VGAT/VGLUT2 mRNAs – containing neurons were clustered around and above the mammillary tract (mt) (Figure 1 A, D) in a region similar to that described in rats as the grandicellular SuM (SuMg) by Pan and Mc Naughton (2004) and that is included in the lateral SuM (SuML) region of Paxinos and Franklin’s mouse Atlas. The numerous singly labeled VGLUT2 mRNA – expressing cells were small and mostly located in the most central area (Figure 1 A) termed the parvicellular SuM (SuMp) by Pan & Mc Naughton (2004) corresponding to the SuMm in Paxinos and Franklin’s mouse Atlas. Some were also intermingled with the larger VGAT/VGLUT2 mRNAs-containing cells in the SuMg. Few neurons expressed VGAT mRNA only. These neurons were scattered within the SuML and SuMM.

**Figure 1:**
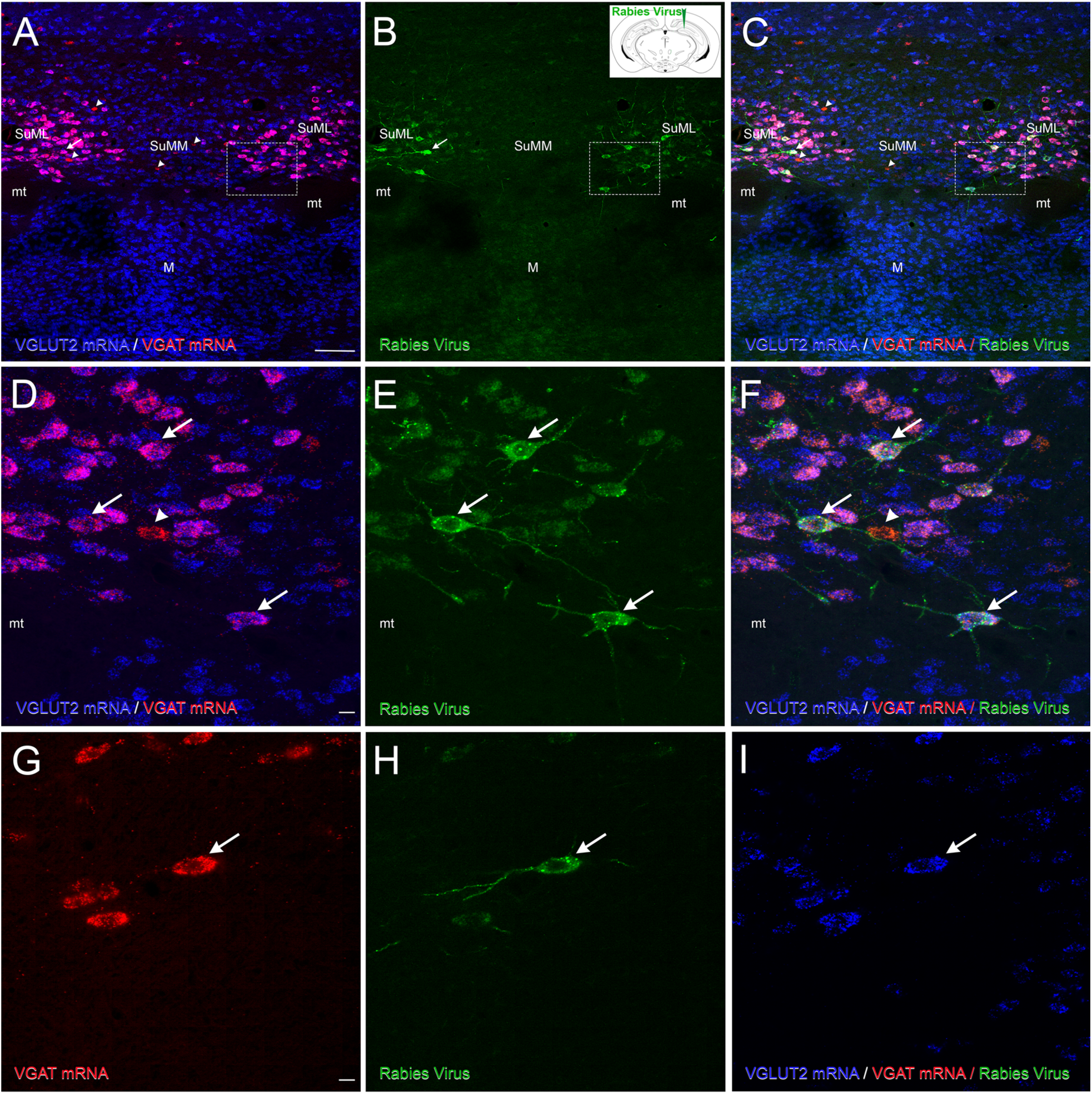
Neurochemical features of SuML neurons innervating the dorsal dentate gyrus (DG) characterized by simultaneous labeling for the rabies virus (RV) retrograde tracer (green), VGAT mRNA (red) and VGLUT2 mRNA (blue) in a coronal section. (A-I) All confocal images were obtained from sequential acquisition of separate wavelength channels, corresponding to the different fluorophores used for the triple labeling, from a single optical slice. This optical slice was acquired in the supramammillary region of the hypothalamus from a coronal section of a VGLUT2-cre mouse that received an injection of RV in supragranular layer of the DG (B, insert). A) Image obtained from the merge of the two confocal images corresponding respectively to the labeling of VGLUT2 (blue) and VGAT (red) mRNAs. Many neurons expressing both VGLUT2 and VGAT mRNAs (arrow) were observed in the lateral region of the supramammillary nucleus (SuML). They were located almost exclusively above and around the mammillary tract (mt). Neurons expressing VGLUT2 mRNA only (blue) were observed mainly in the most medial part of the SuM (SuMM) and were numerous in the mammillary nucleus (M). Few neurons containing VGAT mRNA only (red, arrowheads) were distributed in the SuML and SUMM. B) Confocal image corresponding to the immunohistochemical labeling of the RV (green) showing that all RV containing neurons in the SuM were located in the region of the SuML surrounded the mt. C) Merge of A-B. (D-F) Higher magnification of region outlined in A-C showing that all these SuML neurons projecting to the DG, labeled for the RV (E, arrows), co-expressed VGLUT2 and VGAT mRNAs (D, F, arrows). (G-I) RV labeled neuron (H, arrow) co-expressed VGAT mRNA (G, arrow) and VGLUT2 mRNA (I, arrow) Scale bars: A-C, 100µm; D-I, 10µm.

In summary, our results indicate that in mice like in rats a large number of neurons co-expressing markers of GABAergic and glutamatergic transmission are located in the SuML region.

### Distribution of SuM neurons with dorsal dentate gyrus (dDG) projections

Unilateral or bilateral injections of rabies-virus (RV) were performed into the DG of the dorsal hippocampal formation (Figure 1 B, insert) to evaluate the distribution and proportion of SuM neurons that project to this structure. In all these mice, the RV tracer injection was located either in the granule cell layer or inner molecular region of the dorsal blade of the DG. Within the SuM, all these animals displayed retrogradely labeled neurons exclusively in the region of SuML (SuMg) immediately dorsal to the mt (Figure 1 B, E, H). Virtually none were found in the SuMM (Figure 1 B). Many of these neurons displayed a large soma with several labeled proximal dendrites (Figure 1 E). Therefore in mice, SuM neurons innervating the dDG are located in the SuML as in rats (Soussi et al., 2010).

### Neurotransmitter phenotype of SuM neurons projecting to the dDG revealed by simultaneous detection of RV retrograde tracer, VGAT mRNA, and VGLUT2 mRNA

We next investigated whether the SuML neurons projecting to the dDG express markers of GABAergic and glutamatergic transmissions. Sections processed for simultaneous detection of VGAT mRNA, VGLUT2 mRNA and RV (Figure 1 A-I) showed that almost all RV retrogradely-labeled neurons co-express VGAT and VGLUT2 mRNAs (Figure 1 A-I, arrows). These data were confirmed by quantitative analysis performed on twelve sections (4 mice; 3 sections per mouse) showing that 99% (n= 378 out of 380 neurons) of the RV-labeled neurons co-expressed VGAT and VGLUT2 mRNAs. The percentages of these triple-labeled cells were the same (99%) for the three antero-posterior levels of the SuM analyzed (range 98%-100%).

### Distribution and neurotransmitter phenotype of fibers and axon terminals originating from SuM neurons innervating the dorsal dentate gyrus

To further characterize the neurochemical properties of SuML neurons innervating the dDG, AAV-5-DIO-EYFP was injected bilaterally in the SuML (Figure 2 A) allowing specific EYFP labeling of VGLUT2 neurons and of their axon fibers innervating the hippocampus including the dDG (Figure 2 A). Sections of these VGLUT2-EYFP mice were processed for simultaneous immunohistofluorescent detection of EYFP, VGLUT2 and VGAT or GAD65. All mice injected in the SuML displayed numerous anterograde– EYFP labeled fibers and axon terminals in the dDG (Figure 2 B, arrowheads) on both sides of the hippocampal formation. Labeled fibers and axon terminals were also present in the CA2 ⁄CA3a region of the hippocampus in particular when the injection sites involved more posterior levels of the SuML (Figure 2 B, arrowheads).

**Figure 2:**
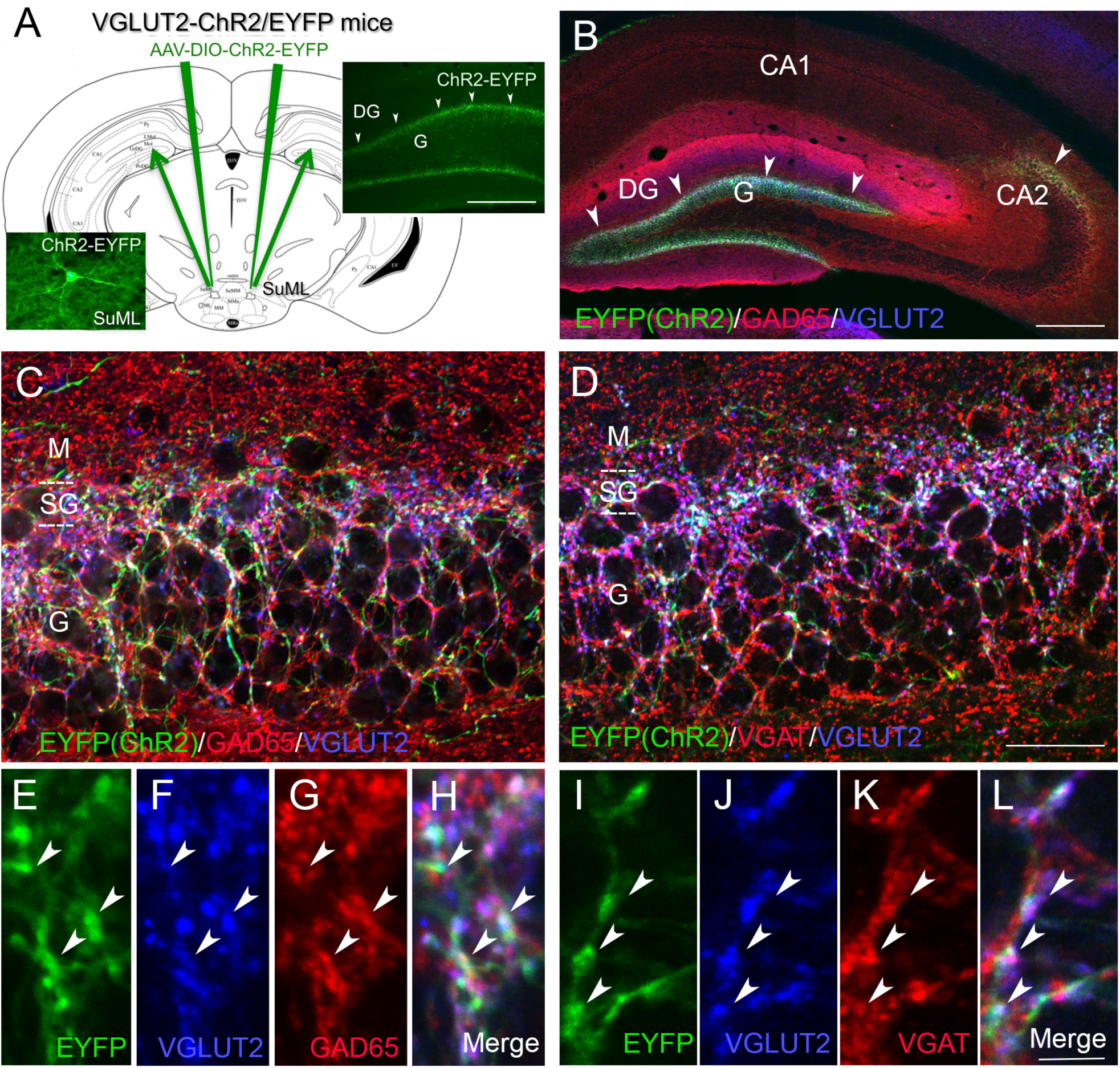
Neurochemical features of axon terminals from SuML neurons innervating the dorsal DG characterized by simultaneous immunohistochemical labeling for the AAV-EYFP anterograde tracer (green), GAD65 or VGAT (red) and VGLUT2 (blue) in coronal sections. (A) Diagram illustrating the bilateral injections of the viral vector AAV-DIO-ChR2-EYFP within the lateral region of the SuM. Images from a coronal section showing endogenous fluorescence of EYFP observed in the cell body and proximal dendrites of a transfected neuron located within the SuML as well as fibers and axon terminals within the supragranular (arrowheads) and granular layer (G) of the DG. (B) Image obtained from a single optical slice showing labeling for EYFP (green), GAD65 (red) and VGLUT2 (blue) in the hippocampus. Fibers labeled with the anterograde tracer AAV-DIO-ChR2-EYFP injected within the SUM as illustrate in (A) were exclusively located in the supragranular (arrowheads) and granule cell (G) layers as well as the CA2 region of the hippocampus. (C, D) Images corresponding to a maximum intensity z-projection of a stack of 8 optical slices spaced at 370 nm, showing labeling for EYFP (green), VGLUT2 (blue) and GAD65 (C, red) or VGAT (D, red) in the dorsal DG. Axon terminals and fibers, from neurons in the SuML, labeled for the EYFP anterograde tracer (green) were located mainly in the supragranular layer but also in the granule cell layer. Numerous GAD65-(C) or VGAT-(D) containing terminals were present in the molecular layer (M) and granule cell layer (G) of the dorsal DG. VGLUT2-containing terminals were mainly located in the supragranular layer (SG) but were also observed in G. (E–L) Images of the three different fluorophores used for the triple labeling, obtained by sequential acquisition of separate wavelength channels from a single optical slice, in the SG of the DG demonstrated that many if not all axon terminals labeled for EYFP (B, I, green, arrowheads) contained GAD65 (G, red arrowheads), VGAT (K, red, arrowheads) but also VGLUT2 (F, J blue, arrowheads). (H) Merge of E-G. (L) Merge of I-K. Scale bars: A-B, 200µm; C-D, 25µm and E-L, 3µm.

In the dDG, EYFP axonal fibers and terminals were mainly located in the supragranular layer of the dorsal and ventral blades of the DG (Figure 2 B-D) although some were localized in the granule cell layers. In all triple labeled sections either for EYFP, GAD65 and VGLUT2 or EYFP, VGAT and VGLUT2, the vast majority if not all EYFP-containing axon terminals present in the supragranular and granule cell layers were labeled for both GAD65 and VGLUT2 (Figure 2 C, E-H) or VGAT and VGLUT2 (Figure 2 D, I-L). Quantitative analysis of double labeling for VGLUT2 and VGAT showed that 90% (range 82% to 99%) of VGLUT2 labeled terminals were labeled for VGAT with no major differences between the infragranular blade (89%; range 82% to 99%) and the supragranular blade (91%; range 83% to 99%). Further quantification of EYFP labeling revealed that 98 % (range 96% and 100%) of the axon terminals contained both VGAT and VGLUT2 confirming that all EYFP-containing axon terminals originating from SuML neurons contained both markers of GABA and glutamate neurotransmissions.

### Electron microscopy analysis of synaptic contacts established by EYFP-containing axon terminals originating from SuML neurons

In the supragranular region of the dDG, axon terminals from SuML neurons labeled for EYFP displayed diffuse electron-dense labeling that contrasted with adjacent unlabeled cellular compartments. These labeled axon terminals formed synaptic contacts on the soma (Figure 3 A, B, D, E, F, I) and dendritic profiles (Figure 3 C, G, H) of presumed granule cells (GCs). These axon terminals were often very large boutons that displayed one (Figure 3 A, B, D, E) or more (Figure 3 F) synaptic zones with relatively thin post-synaptic densities (arrowheads) characteristic of symmetric synapses. Other synaptic contacts formed by these labeled terminals displayed thicker post-synaptic densities characteristic of asymmetric synapses (Figure 3 A, B, C; arrow). We also observed large labeled “en passant” boutons establishing symmetric synapses on the dendrites of a presumed granule cell (Figure 3 G, H). Finally, some axon terminals from SuML neurons formed both symmetric (arrowhead) and asymmetric (arrow) synapses on the soma or the proximal dendrite of two different neighboring GCs (Figure 3, A, B).

**Figure 3:**
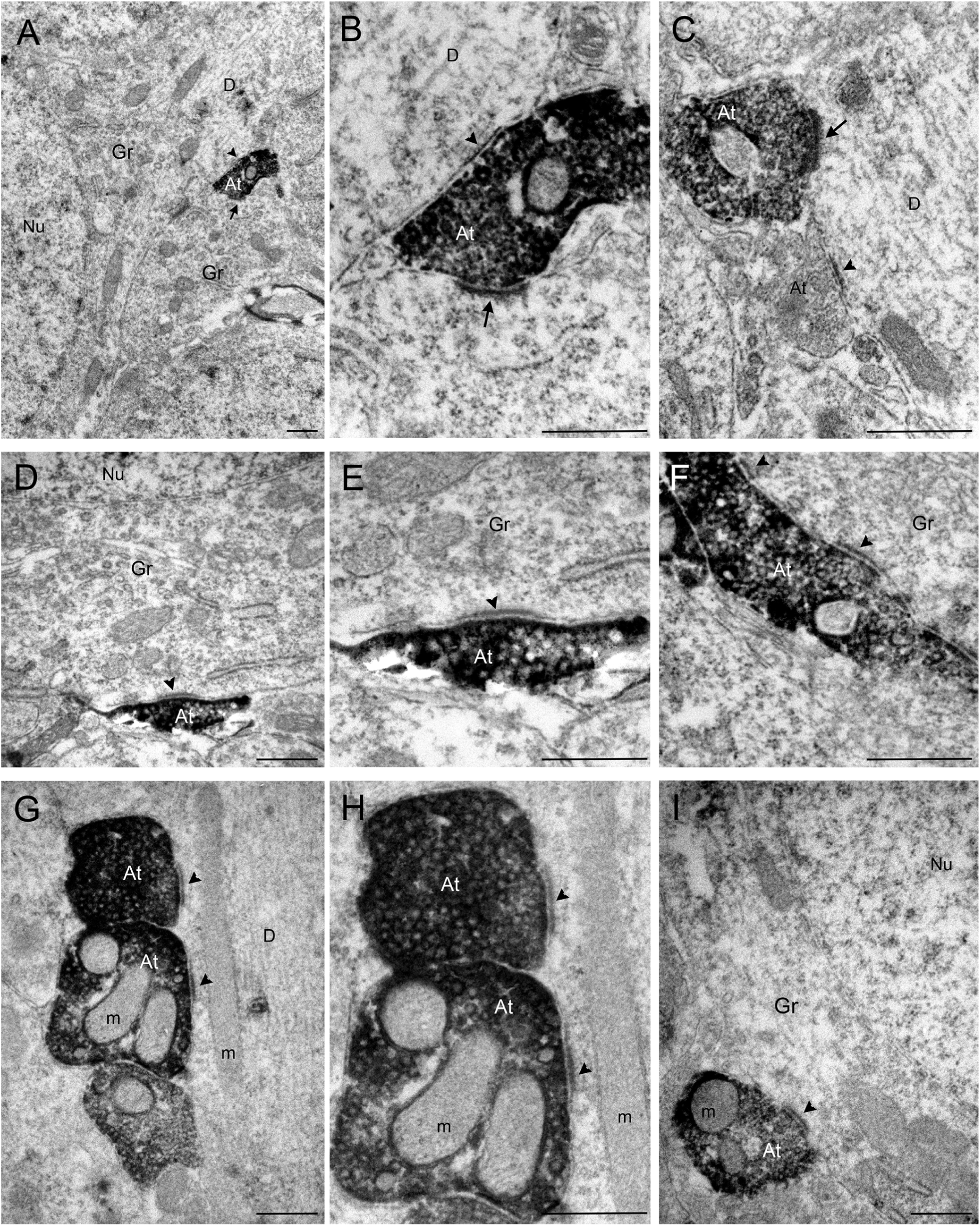
Pre-embedding immunolabeling for EYFP anterograde tracer in ultrathin coronal sections of the dorsal DG. (A–I) In the dorsal DG, numerous axon terminals were labeled for EYFP, revealed by electron-dense peroxidase 3.3’-diaminobenzidine tetrahydrochloride product. (A) A labeled axon terminal (At) making two synaptic contacts (arrow and arrowhead) on unlabeled somata of 2 presumed granule cells (Gr). (B) Higher magnification of the At illustrated in (A) showing that these two synaptic contacts were different: one displayed a relatively thin post-synaptic densities (arrowhead) characteristic of symmetric synapses on the soma of one Gr; the other one displayed a thick post-synaptic densities (arrow) characteristic of asymmetric synapses on the soma of another Gr. (C) Labeled axon terminals established synaptic contacts displaying a thick synaptic densities on unlabeled dendritic processes (D) that differed from the synaptic contact formed by the unlabeled At (arrow). The synaptic contacts illustrated in D-I displayed relatively thin post-synaptic densities (arrowheads) on the soma (D-F, I) or dendrites (G, H) of unlabeled Gr. m, mitochondria; Nu, nucleus. Scale bar: A-I, 0.5 µm.

Together these data demonstrate that in mice all SuM neurons innervating the dDG belong to a single population of large neurons located in the SuMg region of the SuML. These cells display a dual neurochemical phenotype for GABA and glutamate neurotransmissions and establish symmetric (presumably inhibitory) and asymmetric (presumably excitatory) synapses on the GCs of the dDG.

### Co-release of glutamate and GABA at the SuML-dentate granule cells synapses

The neurochemical profile of SuML neurons projecting to the dDG strongly suggested that they co-release GABA and glutamate at the SuML-dentate granule cells synapses. To test this hypothesis, we performed patch clamp recordings of dDG granule cells layer using optogenetic stimulation in hippocampal slices obtained from VGLUT2-ChR2 mice (5 mice, 3 sections per mouse, 16 neurons) and VGLUT2-EYFP control mice (3 mice, 3 sections per mouse, 10 neurons) (Figure 4). The light stimulation of SuML-DG fibers and axonal terminals expressing ChR2 (Figure 4 A) evoked a fast inward and a slower outward synaptic current in fourteen out of sixteen neurons recorded at a holding potential of −30 mV (Figure 4 D, F, H). Post-hoc immunodetection of biocytin-filled neurons showed that all these recorded neurons correspond to GCs (Figure 4 C, E, G, I). They were equally distributed in the apex and the upper and lower blades of dDG granule cell layer (Figure 4 B). These GCs were additionally recorded at different holding potentials. At −10 mV and +10 mV (close to the reversal potential of the glutamatergic receptor-mediated currents), only the outward (positive going) synaptic current was observed (Figure 4 D, see also F, H). From −70 mV to −50mV (close to the reversal potential of GABA-A receptor-mediated currents) only the inward (negative going) synaptic current was observed (Figure 4 D, see also 4 F, 4 H). At intermediate holdings, from −30 to −20mV, the light-evoked synaptic currents displayed both inward and outward components (Figure 4 D, F, H). Using a pharmacological approach, we confirmed the nature of these currents. Bath application of a mixture of AMPA/Kainate and NMDA receptor antagonists (NBQX 10 µM and D-AP5 40 µM) abolished the inward component (Figure 4 F, K, L, red trace) while GABA-A receptor blockers inhibited the outward component (Figure 4 H, J-gabazine; K, L– bicuculline, green trace). Quantitative analysis performed in 5 DG neurons (Fig. 4 K) further illustrated that in regular ACSF the repetitive light pulses (5 ms, 0.05 Hz) evoked postsynaptic currents (PSC) of relatively stable amplitude (1.00 ± 0.26 normalized, Fig. 4 K, L, left hand side red trace is an example of an averaged response of one neuron). The application of glutamate receptor blockers (10 µM NBQX + 40 µM D-AP5) reduced the peak amplitude by 84% (0.16 ± 0.15, p<0.01, n= 5, paired Wilcoxon test). The remaining response seen in 4 out of 5 neurons was probably due to GABA-A mediated current because at −70 mV the driving force for chloride driven currents is close to but is not zero (Figure 4 K, L, red trance in the middle). Therefore in case of GABA massive release some inward current is still possible. Indeed a switch to Vh= 10 mV revealed a large PSC response to light stimulation (Figure 4 L, green trace in the middle) which had a stable amplitude (1.00 ± 0.20 normalized), and was subsquently reduced to 12% (0.12 ± 0.16, p<0.01, n= 5, paired Wilcoxon test) by the addition of 10 µM bicuculline to the ACSF already containing GluR blockers (Figure 4 L, green trace at left). In 3 out of 5 neurons, bicuculline completely abolished the synaptic response to light pulses after 6 min of drug perfusion. In two neurons the small remaining current was probably due to the fact that bicuculline is a competitive antagonist and can be displaced when large quantities of GABA are released. All these results indicate that the inward synaptic current component is mediated by glutamate and the outward component by GABA. Importantly, the disappearance of the inward (glutamatergic) component induced by the light stimulation in the presence of NBQX and D-AP5 antagonists (red trace) did not affect the outgoing (GABAergic) currents (Fig. 4 F). In two GCs (Figure 4 I), the disappearance of the glutamate response (incoming current recorded at −30 mV) in the presence of NBQX and AP5 (red trace) unmasked the GABAergic component (outgoing current) abolished after addition of gabazine (green trace) (Figure 4 J).

**Figure 4:**
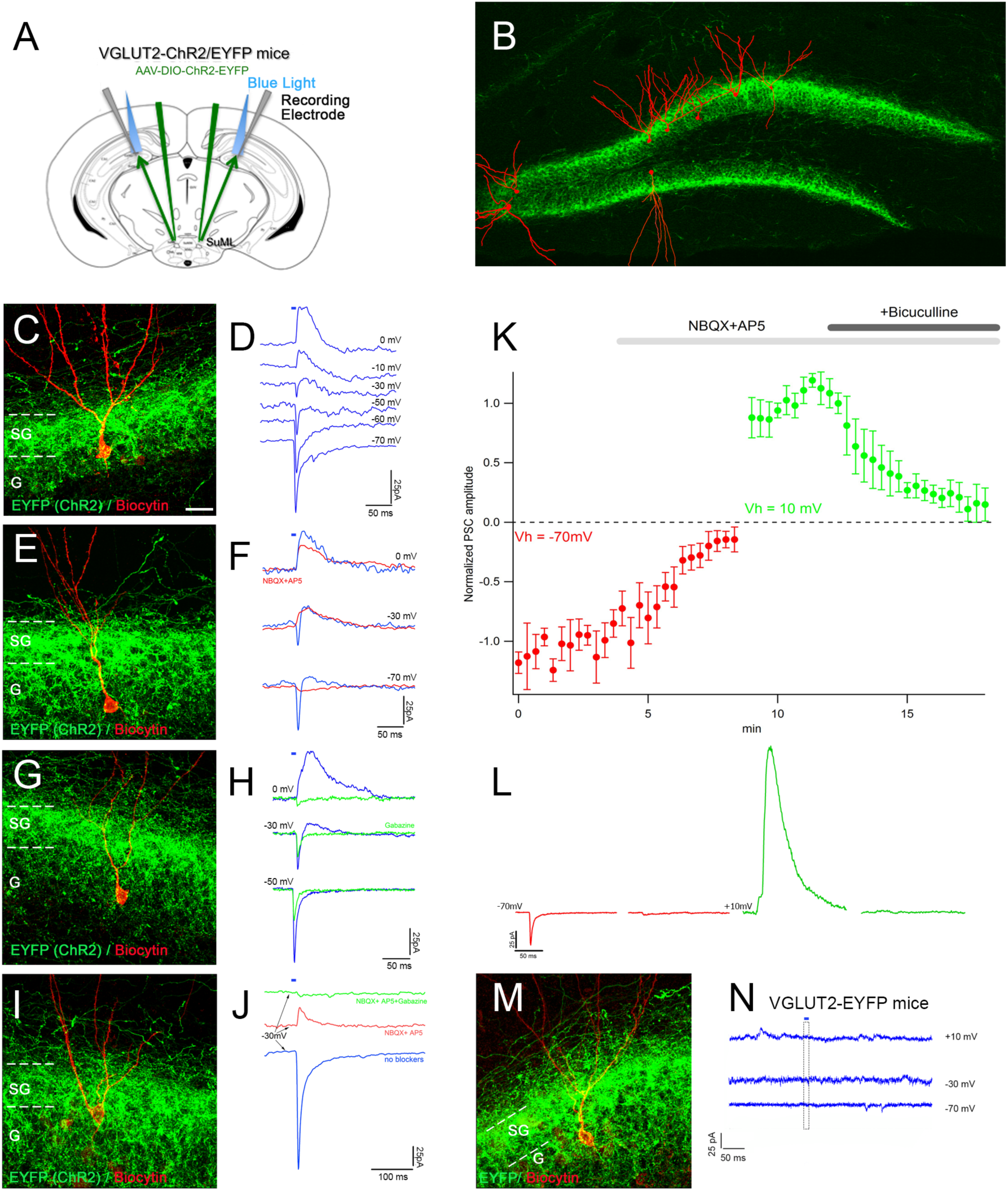
Selective stimulation of axonal terminals from SuML neurons innervating the dorsal DG, performed in hippocampal slices of VGLUT2-ChR2-EYFP mice, induced co-release of GABA and glutamate on DG granule cells. (A) Diagram illustrating the site of the bilateral injections of AAV-DIO-ChR2-EYFP in VGLUT2-cre mice, of light stimulations and of the recorded patch clamp electrode in the DG of the VGLUT2-ChR2-EYFP mice. (B) Montage illustrating the position within the dentate granule cell layer of the biocytin-filled cells reconstructed after patch clamp recordings and EYFP-labeled terminals originating from SuML neurons. (D, F, H, J, N) Examples of light induced PSCs recorded in DG cells illustrated in (C, E, G, I, M). (D) Light induced post-synaptic currents (PSCs) recorded at different holding potentials in DG neuron; short black bar above the upper trace shows the time moment and duration of light stimulus. Note at −70 mV to −50 mV holdings the PSCs had negative-going direction (inward) currents. They were positive-going (outward) at −10 mV and 0 mV but displayed both negative- and positive-going phases when the neuron membrane was clamped at −30 mV. This suggests that light stimulation induces two types of postsynaptic currents. Indeed, the application of glutamate receptor blockers NBQX and AP5 (F, red traces) inhibited inward (F, negative going) component of PSCs recorded at negative holding potentials (Vh), but had only small effect on outward component (positive going) recorded at positive Vh. Inversely, the outward component of light induced PSCs was sensitive to GABA A receptor inhibitor gabazine (H, green vs blue traces). This suggests that these PSCs are generated by simultaneous activation of glutamate and GABA post-synaptic receptors. Interestingly at −50 mV holding, PSC was larger without gabazine (H, blue versus green trace), probably because GABA-mediated current hyperpolarizes the membrane potential increasing thus the driving force for the glutamatergic current. In voltage-clamp condition such an interaction is possible if the pool of postsynaptic GABA Rs is close to the postsynaptic Glu R pool. (K, L) Effect of glutamate and GABA A receptors blockers on the peak amplitude of light pulse evoked PSC recorded at −70 mV (red circles) and 10 mV (green circles) holding potentials. For each Vh the peak amplitudes of first 10 PSCs were averaged and used then as a normalization factor for all peak amplitude recorded at given potential. Each point and error bars corresponds the mean ± SD of PSC normalized amplitude recorded in 5 DG neurons. In regular ACSF the repetitive light pulses (5 ms, 0.05 Hz) evoked PCS of relatively stable amplitude (1.00 ± 0.26, L left hand side red trace-an example of averaged response of one neuron). The application of glutamate receptor blockers (10 µM NBQX + 40 µM D-AP5) reduced the peak amplitude by 84% (0.16 ± 0.15, p<0.01, n= 5, paired Wilcoxon test). The remaining response was seen in 4 from 5 neurons and is probably due to GABA A mediated current because at −70 mV the driving force for chloride driven currents is close to but not zero (L, red trance in the middle) therefore in case of GABA massive release some inward current is still possible. Indeed a switch to Vh= 10 mV revealed a huge PSC response to light stimulation (L, green trace in the middle) which amplitude was stable (1.00 ± 0.20) but progressively reduced to 12% (0.12 ± 0.16, p<0.01, n= 5, paired Wilcoxon test) by the addition of 10 µM bicuculline to the ACSF already containing GluR blockers (L, green trace at left). In 3 from 5 neurons, 6 min lasting bicuculline application completely abolished response to light pulses. In two neurons the remaining current is probably due to the competitive character of bicuculline induced inhibition i.e in case of high GABA release 10 µM of bicuculline may be not sufficient to all receptors inhibition. (M, N) The identical light stimulation (5 ms, 10%, 50% 90% max power LED) of fibers and axon terminals expressing EYFP on slices of control VGLUT2-EYFP mice (n= 3) did not evoke any response in the recorded granule cells (n= 5) confirming that ChR2 activation is required to obtain the PSCs. Scale Bar: C, E, G, I, M, 20µm.

The very short latency of the glutamatergic inward and GABAergic outward currents recorded in a GC after light stimulation of ChR2 containing axon terminals, and the persistence of GABAergic outflow current in the presence of NBQX and AP5 indicates that the GABAergic current results from direct light stimulated GABA release and not from the excitation of DG GABAergic neurons by light stimulated glutamate release. Together, these results suggest that activation of SuML axon terminals produce monosynaptic glutamatergic and GABAergic currents on their targets.

Finally, we verified that light stimulation (5 ms, 10%, 50% 90% max power LED) of fibers and axon terminals expressing EYFP on slices of control VGLUT2-EYFP mice (n= 3) did not evoke any response in the recorded GCs (n= 10) (Figure 4 M, N).

### Effect of optogenetic stimulation of SuML axon terminals innervating the dDG on behavior, LFP and EEG spectral content

We analysed the effects of light stimulation of SuML axon terminals innervating the dDG on behavioral states, spectral content of the LFP recorded in dDG and cortical EEG in VGLUT2-ChR2 (n= 4) and control VGLUT2-EYFP mice (n= 4).

Light activation of SuML axon terminals in VGLUT2-ChR2 mice during waking (WK), induced a strong and significant increase in EMG value as compared to control mice (mean EMG ratio, VGLUT2-EYFP= 0.9 ± 0.03, VGLUT2-ChR2= 2.0 ± 0.31, p= 0.0209). The increase in EMG was due to an increase in animal movements (Figure 5 A, D). It was associated with a slight but not significant increase in theta power in the dDG LFP (Figures 5 B; 6 A) and the EEG (Figure 7 A) compared to control VGLUT2-EYFP mice. However, theta/delta ratio was significantly increased both in the dDG LFP (Figure 6 A, B) and the EEG (Figure 7 C, D) during the stimulation compared to control mice (LFP: VGLUT2-ChR2= 1.3 ± 0.11, VGLUT2-EYFP= 0.9 ± 0.02, p=0.0209, EEG: VGLUT2-ChR2= 2.2 ± 0.31, VGLUT2-EYFP= 0.9 ± 0.05, p=0.0209). The frequency at the theta peak was slightly but not significantly increased during stimulation as compared to control VGLUT2-EYFP animals (LFP: VGLUT2-ChR2: 8.43 ± 0.43 Hz, VGLUT2-EYFP: 6.96 ± 0.23 Hz, p=0.0833) (Figures 5 B; 6 D). Gamma power (30-100 Hz) was significantly increased during the stimulation in VGLUT2-ChR2 mice as compared to control mice in the dDG LFP (VGLUT2-ChR2= 1.6 ± 0.13, VGLUT2-EYFP= 1.0 ± 0.02, p=0.0209) (Figures 5 C; 6 A, C) and in the EEG (VGLUT2-ChR2: 1.68 ± 0.05, VGLUT2-EYFP: 0.98 ± 0.02, p=0.0209 (Figure 7 B, C, E). The increase occurred immediately after the beginning of the stimulation and lasted until the end of it (Figures 6 C; 7 E).

**Figure 5:**
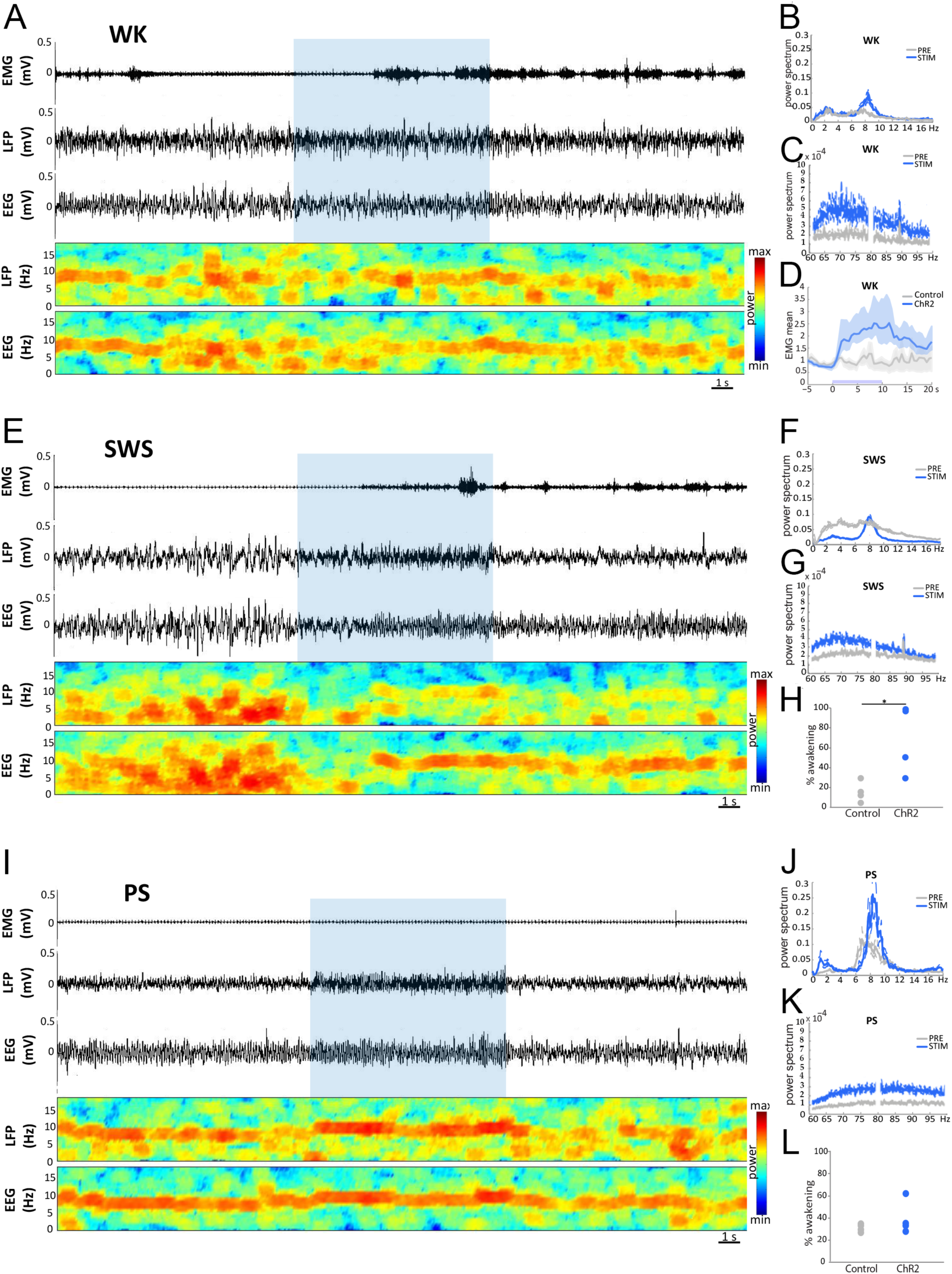
Effects of light stimulation of axonal terminals from SuML neurons innervating the dorsal DG on DG LFP, EEG recordings and behavior of VGLUT2-ChR2-EYFP mice during WK, SWS and PS. (A, E, I) Examples of raw recording for EMG, LFP recorded in the DG and parietal cortex EEG as well as associated time-frequency analysis of LFP and EEG in a VGLUT2-ChR2-EYFP mouse during WK (A), SWS (E) and PS (I). The blue bar represents the optogenetic stimulation period (20 Hz with pulses of 10 ms for 10 s). (B, C, F, G, J, K) LFP power spectra between 0-18 Hz and 60-100 Hz in VGLUT2-ChR2-EYFP mouse before and during light stimulation (20 Hz 10 s, pulses of 10 ms, during 4 h) during WK (B, C), SWS (F, G) and PS (J, K). An effect was clearly visible on the LFP in all vigilance states. The stimulation induced a slight increase in the theta power during WK (A, B) and a major increase in theta power and frequency during PS (I, J) as well as a clear reduction of slow waves oscillation during SWS (E, F). The stimulation during WK and PS also increased the power of gamma (C, K). Light stimulation during WK increased locomotor activity reflected by an increase of EMG signal in VGLUT2-ChR2-EYFP mice (A, D, blue trace) as compared to control VGLUT2-EYFP mice (D, grey trace). (H) Light stimulation during SWS induced awakening reflected by a significant increase of the awakening percentage after stimulation in most VGLUT2-ChR2-EYFP mice (range 30% to 100%) as compared to that observed in control VGLUT2-EYFP mice (range: 0% to 30 %). (L) In contrast no significant difference in the percentage of awakening was observed between VGLUT2-CHR2-EYFP and control VGLUT2-EYFP mice when stimulation was performed during PS.

**Figure 6:**
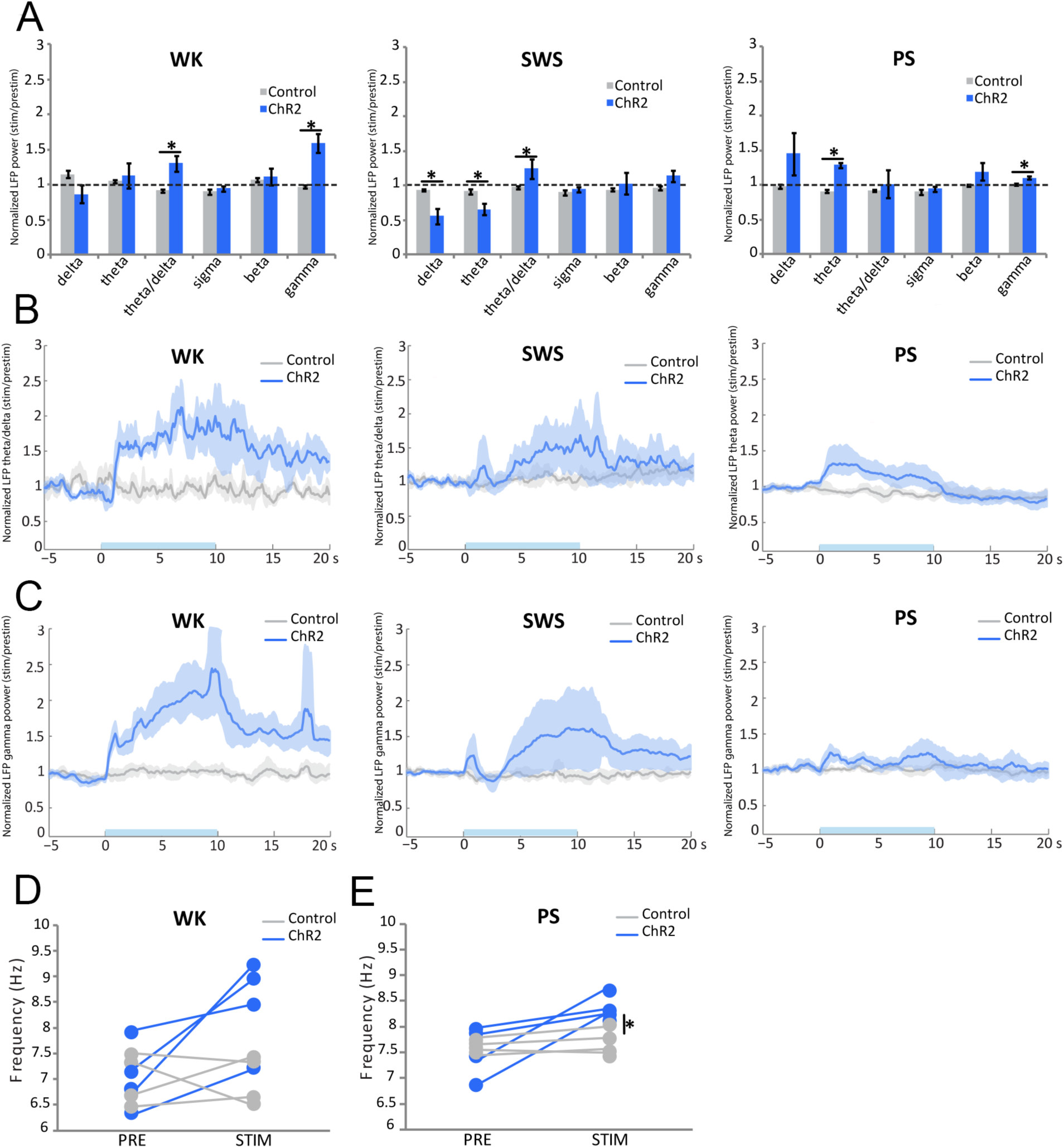
Comparison of the effects of light stimulation of axonal terminals from SuML neurons innervating the dorsal DG on DG LFP performed during WK, SWS and PS between VGLUT2-ChR2-EYFP and control VGLUT2-ChR2-EYFP mice. (A) Ratio of power recorded during light stimulation to that recorded before the stimulation (20 Hz, 10 s for 4 h with pulses 10 ms) for the different frequency bands (delta, theta, gamma) of LFP in VGLUT2-ChR2-EYFP (n= 4) and control VGLUT2-EYFP (n= 4) mouse groups during WK, SWS and PS. These ratios illustrate: an increased power in the theta frequency band during stimulations performed in PS; an increase of theta/delta power during the stimulation when performed in WK and SWS and an increased power in the gamma frequency band when stimulations were performed during WK and PS in the VGLUT2-ChR2-EYFP group. Increased power observed during stimulations observed in the ChR2 group differed significantly from random variation observed before and after the stimulation in the control group. Significance: Mann Whitney: * p <0.05. (B, C) Theta/delta (B) and gamma (C) power ratios as a function of time after light stimulation during the different vigilance states. The blue bar on the x axis represents the stimulation period. Shaded regions show 95% confidence intervals. During WK, a significant increase of theta/delta (B) and gamma (C) power occurred in VGLUT2-ChR2-EYFP group immediately after the light stimulation. These increases were observed during the entire 20 s period analyzed. The increase of theta/delta during SWS occurred two to three seconds after the stimulation. The increase of theta (B) and gamma (C) powers during PS occurred immediately after the stimulation and stayed during the 10 s period of stimulation. No difference induced by the stimulation was observed in the control group. (D, E) Peak frequency analysis of theta before and during optogenetic stimulation in the control group (n=4) and the ChR2 group (n= 4). Significance: Mann Whitney, * p <0.05 compared to the control group.

**Figure 7:**
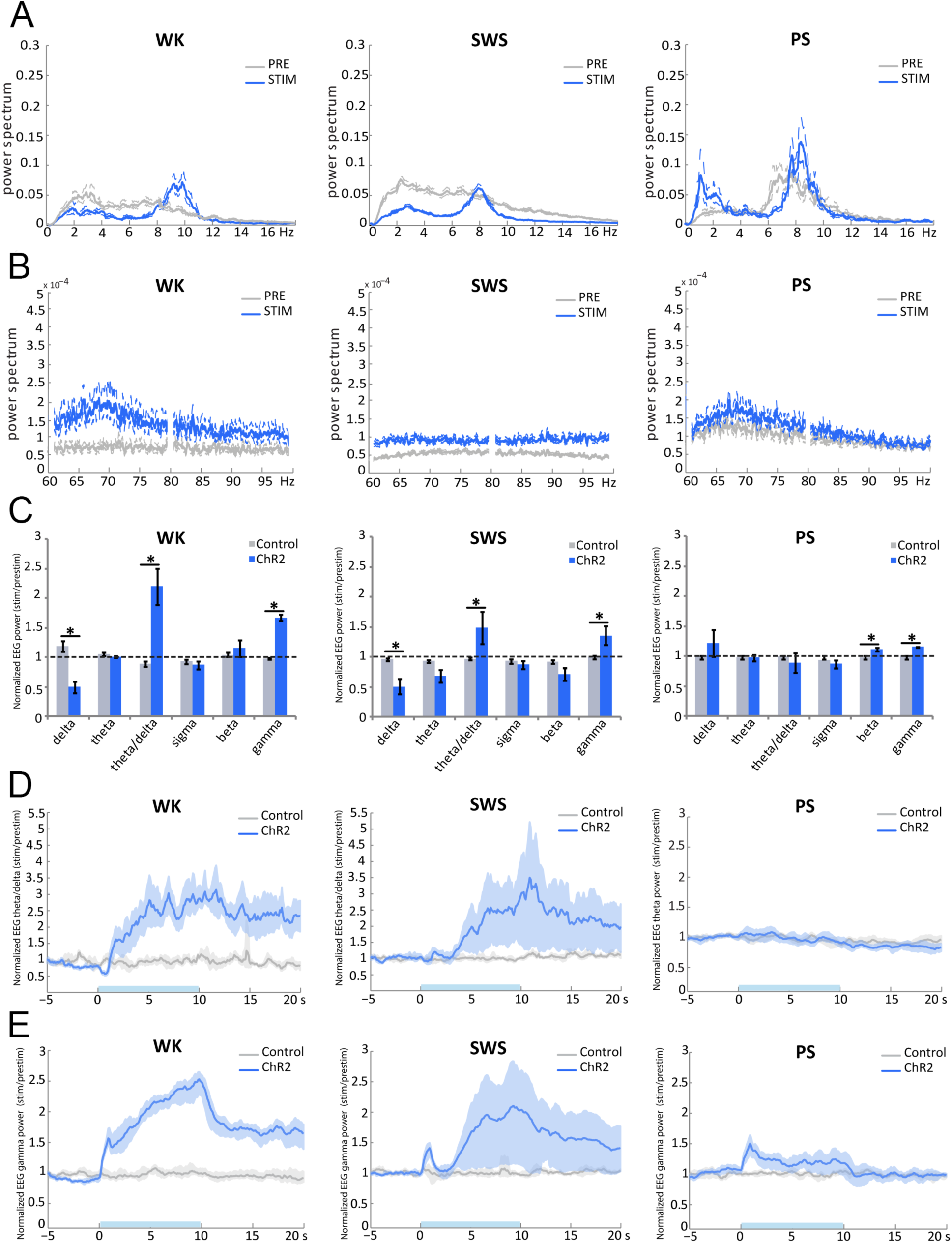
Comparison of the effects of light stimulation of axonal terminals from SuML neurons innervating the dorsal DG on EEG performed during WK, SWS and PS between VGLUT2-ChR2-EYFP and control VGLUT2-EYFP mice. (A, B) EEG power spectra between 0-18 Hz and 60-100 Hz in a VGLUT2-ChR2-EYFP mouse before and during light stimulation (20 Hz 10 s, pulses of 10 ms, during 4 h) during WK, SWS and PS. An effect was clearly visible on the EEG in all vigilance states. (C) Ratio of power recorded during light stimulation to that recorded before the stimulation for the different frequency bands (delta, theta, gamma) of EEG in VGLUT2-ChR2-EYFP (n= 4) and control VGLUT2-EYFP (n= 4). These ratios illustrate an increase of theta/delta power during the stimulation when performed in WK and SWS and an increased power in the gamma frequency band when stimulations were performed during all vigilance states in the VGLUT2-ChR2-EYFP group. These increased powers during stimulations observed in ChR2 group differed significantly from random variation observed before and after the stimulation in the control group. Significance: Mann Whitney: * p <0.05. (D, E) Theta/delta (D) and gamma (E) power ratios as a function of time after light stimulation during the different vigilance states. The blue bar on the x axis represents the stimulation period. Shaded regions show 95% confidence intervals. During WK, a significant increase of theta/delta (B) and gamma (C) powers occurred in VGLUT2-ChR2-EYFP group immediately after the light stimulation. These increases were observed during the entire 20 s period analyzed. The increase of theta/delta during SWS occurred two to three seconds after the stimulation. The increase of gamma (C) powers during PS occurred immediately after the stimulation and lasted during the 10 s period of stimulation. No increase in theta was observed during PS (C, D) in the EEG. No difference induced by the stimulation was observed in the control group.

Stimulations during SWS induced significantly more often an awakening in VGLUT2-ChR2 than in control VGLUT2-EYFP mice (p=0.0209, VGLUT2-ChR2: 80 % of the cases, range 40%-100%; VGLUT2-EYFP: 15 % of the cases, range: 0%-30%; Figure 5 E, H). The induced waking state lasted at least the duration of the stimulation and was characterized by a significant increase in theta/delta ratio (LFP: VGLUT2-ChR2= 1.3 ± 0.14, VGLUT2-EYFP= 1.0 ± 0.02, p=0.0433; EEG: VGLUT2-ChR2= 1.5 ± 0.27, VGLUT2-EYFP= 1.0 ± 0.03, p=0.0209) (Figures 6 A, B; 7 C, D) and a decrease of delta power both in the dDG LFP (VGLUT2-ChR2= 0.6 ± 0.11, VGLUT2-EYFP= 0.9 ± 0.01, p=0.0209) and the EEG (VGLUT2-ChR2= 0.5 ± 0.13, VGLUT2-EYFP= 0.9 ± 0.02, p=0.0209) in VGLUT2-ChR2 compared to control VGLUT2-EYFP mice (Figures 5 E, F; 6 A; 7 A, C). This induced waking state was also associated with an increase in gamma power in the dDG LFP (Figures 5 E, G; 6 A, C) and in the EEG (VGLUT2-ChR2= 1.36 ± 0.16, VGLUT2-EYFP 0.99 ± 0.03, p=0.0433) (Figure 7 B, C, E).

In contrast to SWS, light stimulation of SuML axon terminals in the dDG of VGLUT2-ChR2 mice during PS did not induce significantly more awakening in VGLUT2-ChR2 mice compared to control VGLUT2-EYFP animals (p= 0.3865; Figure 5 L).

Light activation of SuML axon terminals during PS in VGLUT2-ChR2 induced a significant increase in theta power compared to VGLUT2-EYFP mice in the dDG LFP (Figures 5 I, J; 6 A, B; VGLUT2-ChR2= 1.3 ± 0.04, VGLUT2-EYFP= 0.9 ± 0.02, p=0.209) but not in the EEG (Figure 7 C, D; p= 0.773). The frequency at the theta peak was also significantly increased in the dDG LFP compared to control EYFP-VGLUT2 animals (VGLUT2-ChR2: 8.3 ± 0.13 Hz, VGLUT2-EYFP: 7.7 ± 0.15 Hz, p= 0.0209) (Figure 6 E). In addition, gamma was significantly increased both in the dDG LFP and the EEG during the stimulation in VGLUT2-ChR2 mice compared to control VGLUT2-EYFP mice (LFP: VGLUT2-ChR2= 1.1 ± 0.02, VGLUT2-EYFP= 1.0 ± 0.01, p= 0.0209; EEG: VGLUT2-ChR2: 1.68 ± 0.05, VGLUT2-EYFP: 0.98 ± 0.02, p= 0.0209; Figures 5 K, 6 A, C; 7 B, C, E).

### Effect of optogenetic activation of SumL axon terminals innervating dDG on cFos expression

Mouse brains were processed for immunohistochemical detection of cFos in order to assess the effect of SuML axon terminal activation on DG cell activity. Whereas only a few neurons were labeled for cFos in the GCL of the DG in control VGLUT2-EYFP mice (n= 4) (Figure 8 A, B), numerous neurons strongly labeled for cFos were observed in the granule cell layer of the dDG in VGLUT2-ChR2 mice (n= 4) (Figure 8 C, D). Quantitative analysis showed a 241% increase in the number of labeled cFos neurons in the dDG ipsilateral to the stimulation site (270 ± 33) as compared to control mice (ipsi: 79 ± 23; p= 0.026; Figure 8 E) whereas no significant difference was observed in the contralateral dDG (VGLUT2-ChR2: 178 ± 21; VGLUT2-EYFP contra: 90 ± 33; p=0.2; Figure 8 E).

**Figure 8:**
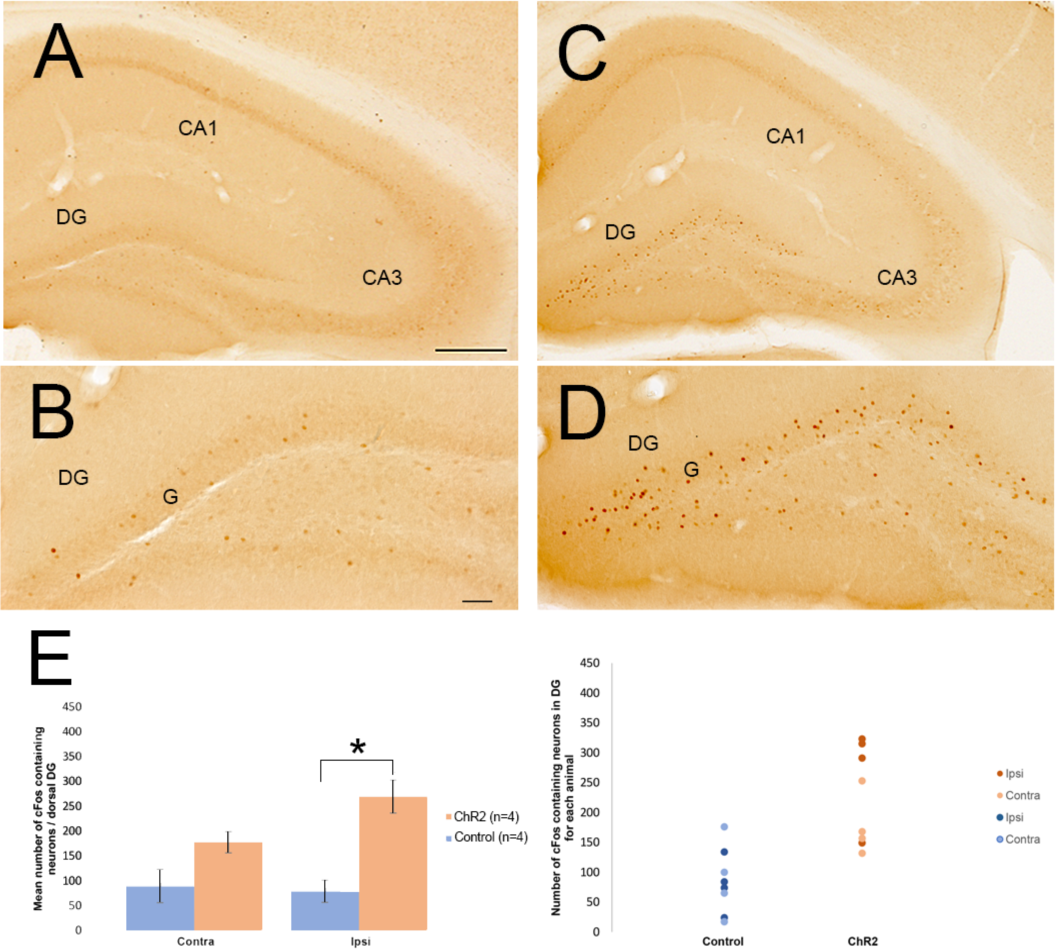
Activation of DG cells after light stimulation of axon terminals from SuML neurons innervating the DG in VGLUT2-ChR2-EYFP and VGLUT2-EYFP as reflected by cFos immunohistochemical labeling. (A, B, C, D) Immunohistochemical labeling for cFos on sections of control VGLUT-EYFP (A, B,) and VGLUT2-ChR2-EYFP (C, D) mice. In control mice (A, B) only very few neurons labeled for cFos were observed in the hippocampus. There were scattered in all layers of the hippocampus including CA1, CA3 and the DG. In addition, these neurons were lightly labeled for cFos including in the granule cell layer (G) of the DG. In contrast, in VGLUT2-ChR2-EYFP mice, many c-Fos containing neurons were located in the granule cells layer (G) of the DG whereas only a few were observed within the CA3 and CA1 regions of the hippocampus. In the granule cell layer these neurons were highly labeled for cFos (D) as compared to those observed in control mice (B). (E) Quantitative analysis confirmed a clear increase number of cFos-containing neurons in the granule cell layer of the DG stimulated (ipsi) but not in the contralateral DG (contra) in VGLUT2-ChR2-EYFP mice (n= 4) as compared to control mice (n= 4). Wilcoxom Rank Sum test * P< 0.05%; Scale Bars: A, C: 250 µm; B, D: 50 µm.

## Discussion

### Neuroanatomical and “in vitro” experiments showing the dual glutamate-GABA feature of the SuML-DG pathway

Our results first establish at the neuroanatomical level that in mice as in rats (Soussi et al., 2010), all SuM neurons innervating the dDG display a dual GABAergic and glutamatergic neurotransmitter phenotype. We further demonstrate that these neurons correspond to a population of SuM cells located just dorsal to the mammillary tract that likely correspond to grandicellular neurons described in rat within the SuML region (Paxinos and Watson 1998) or SuMg (Pan and McNaughton 2004). In addition, our EM data show that SuML terminals form asymmetric (presumed excitatory glutamate) synapses onto some GCs and symmetric (presumed inhibitory GABA) synapses onto others as previously described in rat (Boulland et al., 2009; Soussi et al., 2010). We further found in this study that one axon bouton from a SuML neuron can form an asymmetric synapse on a GC and a symmetric synapse on another GC. In agreement with these neuroanatomical results, our *in vitro* electrophysiological experiments show that optical stimulation of SuM axon terminals innervating the dDG induce co-release of GABA and glutamate on almost all dentate GCs in line with two recent studies (Pedersen et al 2017; Hashimotodani et al., 2018). Hashimotodani et al. (2018) further showed that such co-transmission of GABA and glutamate induce net excitatory effects on GCs and potentiate GC firing when temporally associated with perforant path inputs. In line with such hypothesis, after *in vivo* optic stimulation of SuML axon terminals innervating the dDG in VGLUT2-ChR2 mice, we found that a significant number but not all GCs neurons below the optic fiber were labeled with cFos. Therefore, these cFos labeled neurons could constitute a population of GCs that are simultaneously activated by the optic stimulation of SuML axon terminals and perforant path inputs.

### Effect of optogenetic stimulation of SuML-DG fibers on vigilance states

Activation of SuML axon terminals projecting to the DG induces an awakening effect in mice when performed during SWS but not during PS. It has been shown by Renouard et al. (2015) that the SuML-DG pathway is active during PS and therefore its stimulation during this state might not induce WK because the pathway is already engaged and its overactivation might therefore not to be sufficient to awaken the animal. In contrast, when the stimulation occurs during SWS, an awakening is induced likely because the path is normally inactive during this state. It can be proposed that the stimulation of the DG granule cells by the SuML induces the reactivation of memories and subsequently of structures involved in the exploration leading to an awakening of the animal during SWS. It might also be due to the fact that there is no muscle atonia during this state compared to PS. In line with such hypothesis, stimulation during WK induces increased motor and exploratory activity. Such result is in line with the literature since an increase in exploratory activity was reported when stimulating with blue light dDG neurons expressing ChR2 (Kheirbek et al., 2013).

### Optogenetic stimulation of SuML-DG fibers increases gamma and theta

Our study further demonstrates that activation of SuML axon terminals innervating the DG during PS increases theta power and frequency as well as gamma power in the DG LFP and to a minor extent in the EEG. Activation of the SuML fibers during WK also induces an increase of gamma power in the DG LFP and EEG associated with an increased locomotion. During SWS, the activation of the SuML-dDG pathway induced awakening and a switch from delta to theta activity and an increase in gamma power both in DG LFP and EEG.

It has been previously shown that the SuM can exert significant modulatory control of the theta rhythm (Vertes and Kocsis, 1997). A large percentage of SuM neurons discharge rhythmically, in phase with theta (Kirk and McNaughton, 1991, Kocsis and Vertes, 1994) and this activity is independent of that occurring in the hippocampus. Indeed, neurons in the SuM continue to fire bursts in the theta range frequency after lesion or pharmacological inactivation of the medial septum (Kirk and McNaughton, 1991) known to abolish theta rhythmic activity in the hippocampus. Further, electrical stimulation or carbachol injections in the SuM synchronously drive theta phase-locked cells in both septum (Bland et al., 1994) and hippocampus (Colom et al., 1987). In addition, the SuM controls the frequency and amplitude of theta (Kirk and McNaughton, 1993). These results indicated that the SuM plays a role in theta occurrence but experiments were performed in anesthetized animals and did not specifically study the SuML-DG pathway like in the present study. Indeed, SuM neurons also project directly to the medial septum (Vertes and McKenna, 2000) and can influence theta through the latter structure as well.

Our results also show that the frequency at the peak of theta is higher during PS than in WK in basal conditions. Further, we found out that optical stimulation of SuML-DG fibers during PS induced a similar increase of the frequency at the peak of theta. Besides, lesion of the SuM induced a decreased of theta power specifically during PS (Renouard et al., 2015). On the other hand, lesion of all neurons or specific inactivation of the GABAergic neurons of the medial septum strongly decreases theta during PS and WK (Mitchell et al., 1982, Green and Arduini, 1954, Boyce et al., 2016). Our data also show that stimulation of the SuML-DG fibers induces an increase in gamma power in the dDG and also in the EEG both during PS and WK. Interestingly, Montgomery et al. (2008) found by coherence analysis that dentate/CA3 theta and gamma power and synchrony were significantly higher during PS compared with active WK and that, in contrast, gamma power in CA1 and CA3-CA1 gamma coherence showed significant decreases during PS. These and our results strongly suggest that the medial septum GABAergic neurons induces theta both during WK and PS whereas the increase of theta and gamma power and theta frequency occurring in the DG during PS compared to WK is induced by the projection from the SumL.

### Functional role of the SumL-DG pathway

It has been previously shown that GCs of the DG are instrumental for spatial discrimination (McHugh et al., 2007). In particular, it has been shown that small populations of GCs (2-4%) representing memory engrams are specifically activated when the animals are exposed to a specific context. These cells are reactivated each time the animal is re-exposed to the same context (Schmidt et al., 2012). Optogenetic activation in a different context of a DG engram activated during contextual fear conditioning induces freezing (Liu et al., 2012). Conversely, inactivation during contextual fear memory recall of the DG GCs activated during encoding decreases freezing (Denny et al., 2014). These results indicate that activation of memory engrams composed of DG GCs cells is necessary for spatial learning. Interestingly, in these studies, the mean number of neurons labeled with cFos or Arc in one section (35 µm) of the DG after contextual fear conditioning was in the 30-60 range. In the present study, during stimulation of the SuML fibers in the DG, a mean of 67 DG cells were expressing cFos in the DG per section (40 µm). Finally, Renouard et al. (2015) counted a mean of 68 Arc- and 66 cFos-labeled neurons in one DG section after PS hypersomnia in rats. Therefore, approximatively the same number of DG GCs cells is activated during encoding of a contextual fear memory, stimulation of the SuML fibers in the DG and PS hypersomnia. First, the fact that a similar number of cells are activated when stimulating SuML terminals as during PS hypersomnia suggests that the activation seen during PS is likely due to activation of the SuML-DG pathway. Second, the fact that a similar number of cells are activated during a memory task as during PS hypersomnia and stimulation of the SuML fibers suggest that memory engrams could be activated in these two conditions. Therefore, the induction of an active behavior when the stimulation is made during WK could be due to the activation of memory engrams. Since only a limited number of GCs cells are activated during PS and SuML terminals stimulation despite the fact that a large number of GCs neurons seems to be innervated by SuML axon terminals, it also suggests that activation of these cells is due to a conjunction of the SuML input with another excitatory input. It can be proposed that the medial entorhinal input is involved since it is the main excitatory afferent to the DG involved in its activation during memory encoding (Sasaki et al., 2015). Such hypothesis remains to be tested using optogenetic manipulation of the activated neurons during PS.

## Acknowledgements

This work was supported by INSERM and Aix-Marseille University (M. E, H. E., C. B., L. C., AI. I, A. G., J. S.,); Partenariats Hubert Curien (PHC) IMHOTEP (M.E., H.E., NE. A.); Agence Universitaire Francophone (H.E); CNRS, Fondation pour la recherche médicale (FRM), Societé Francaise de Recherche et Médecine du Sommeil (SFRMS), University Claude Bernard of Lyon (F.B., P-H. L., P-A. L., S. A.) and NS104590 (I. S., E. K-M). We thank the animal facility (CEFOS, AMU, Marseille), the imaging facility (INPHIM, AMU, Marseille) and the electron microscope facility (IBDM, Marseille).

